# Dynamics of cell states and alternative splicing following kidney ischemia-reperfusion injury

**DOI:** 10.1101/2024.10.24.620070

**Authors:** Dana Markiewitz, Jacob Goldberger, Tomer Kalisky

## Abstract

The progression of kidney damage in chronic kidney disease (CKD) involves multiple post-injury stages and complex cellular and molecular mechanisms that are not yet fully understood. In our study we set to characterize the dynamics of mRNA splicing following kidney injury. To this end, we analyzed publicly available bulk RNA-seq data covering nine time points following a kidney ischemia-reperfusion injury (IRI) mouse experiment. Using topic modeling we discerned five distinct temporal phases corresponding to the following cell states: “early injury response”, “injury”, “repairing”, “failed recovery”, and “healthy proximal tubule”. Additionally, we discovered a set of genes that are alternatively spliced between selected time points associated with these cell states, some of which are related to injury, stress, EMT, and apoptosis. Finally, we found several putative splicing regulators that are differentially expressed between the different time points and whose binding motifs are enriched in the vicinity of alternatively spliced exons, indicating that they may play critical roles in mRNA splicing dynamics following kidney injury and repair. These findings enhance our understanding of the molecular mechanisms involved in kidney injury and repair, offering potential avenues for developing targeted therapeutic strategies for acute kidney injury (AKI) and its progression to CKD.

## INTRODUCTION

Acute kidney injury (AKI) is a condition characterized by an abrupt loss of kidney function, and is associated with increased mortality rates, prolonged hospital stays, and elevated healthcare costs (Chertow et al. 2005; Coca, Singanamala, and Parikh 2012; Rewa and Bagshaw 2014). AKI is mainly caused by ischemic or nephrotoxic injury, leading to death of epithelial cells in the nephron accompanied by inflammation and dedifferentiation in surviving epithelia. In order to restore the damage to the nephron, the dedifferentiated epithelial cells proliferate and redifferentiate (Kusaba et al. 2014). This inherent repairing mechanism of the kidney usually allows functional recovery with supportive care over time. However, in cases of persistent residual inflammation and fibrosis, normal kidney function may not be fully restored, thereby increasing the risk of developing chronic kidney disease (CKD) and, potentially, kidney failure (Chawla et al. 2014; Parr and Siew 2016). Despite extensive research, clinical outcomes have not significantly improved over the past decade, and the molecular mechanisms determining successful or failed nephron repair remain unclear.

In recent years, several studies have aimed to characterize kidney damage and repair through the use of single-cell genomics (X. Guo et al. 2022; Kirita et al. 2020; Rudman-Melnick et al. 2020). Kirita et al. (Kirita et al. 2020) used single nucleus RNA sequencing (snRNA-seq) to characterize cell states in a bilateral ischemia-reperfusion injury (IRI) mouse model from 4 hours to 6 weeks following injury. They identified cell clusters representing distinct states of the proximal tubule and labeled them as “severely injured”, “injured”, “repairing”, “healthy”, and “failed repair”. The “severely injured” proximal tubular cells, identified by overexpression of the genes Hspa1a and Krt20, and the “injured” cells, over- expressing the gene Myc and the general kidney injury marker Havcr1 (Kim-1), were present mainly during the first 12 hours following injury. The “repairing” proximal tubular cells, marked by over- expression of the gene Top2a, were predominant 2 days after injury. In contrast, the “failed repair” cell population, marked by the genes Vcam1, Dcdc2a, Sema5a, and Ccl2, was first observed after 2 days and persisted for up to 6 weeks post-injury. A similar study identified a cell population resembling the “failed repair” cells also in the human kidney (Muto et al. 2021). This population had an increased expression and chromatin accessibility for the failed repair marker VCAM1.

Time dependent bulk RNA sequencing measurements were also performed on kidney injury models (Craciun et al. 2016; Liu et al. 2017) in order to identify transcripts associated with different stages of injury and recovery. For example, Liu et al. (Liu et al. 2017) found that genes related to early stress response were elevated for a few hours following IRI, while transcripts related to the cell cycle and wound repair peaked at day 2 after injury. At later times, genes related to inflammation and fibrosis were over-expressed from 24 hours up to 4 weeks following IRI, while those related to normal tubular function were downregulated throughout this period and recovered only after 6 months.

In this study, we set to characterize the dynamics of mRNA splicing following kidney injury. To this end, we used publicly available bulk RNA-seq data collected by Liu et al. (Liu et al. 2017) at different time- points following bilateral kidney IRI in mice, as well as sham surgery and normal controls. We first used topic modeling, an unsupervised machine learning technique, to identify the underlying cell states from the bulk RNAseq expression profiles. This allowed us to represent the series of gene expression measurements as a weighted sum of five cell states with time-dependent coefficients. We characterized these cell states as labeled them as “early injury response”, “injury”, “repairing”, “failed recovery”, and “healthy proximal tubule”. We then identified transcripts that are alternatively spliced between selected time points associated with these cell states, as well as putative splicing regulators. We anticipate that these findings will advance our understanding of the mechanisms of mRNA splicing involved in kidney injury and repair.

## RESULTS

Topic modeling of gene expression levels from a mouse kidney following IRI reveals time-dependent cell states We downloaded an RNA-seq dataset collected by Liu et al. (Liu et al. 2017) containing samples from nine time points, ranging from 2 hours to 12 months following IRI, as well as sham surgery and normal controls. We then performed sequence alignment, obtained a gene expression counts matrix (Table S1), and fitted a topic model with k=5 topics (Figure 1A and Table S2). We identified the cell state associated with each topic using sets of known marker genes that we compiled from the literature (Table S3) and GO enrichment analysis (Figure 1B, Tables S4-S8).

**Figure 1:**
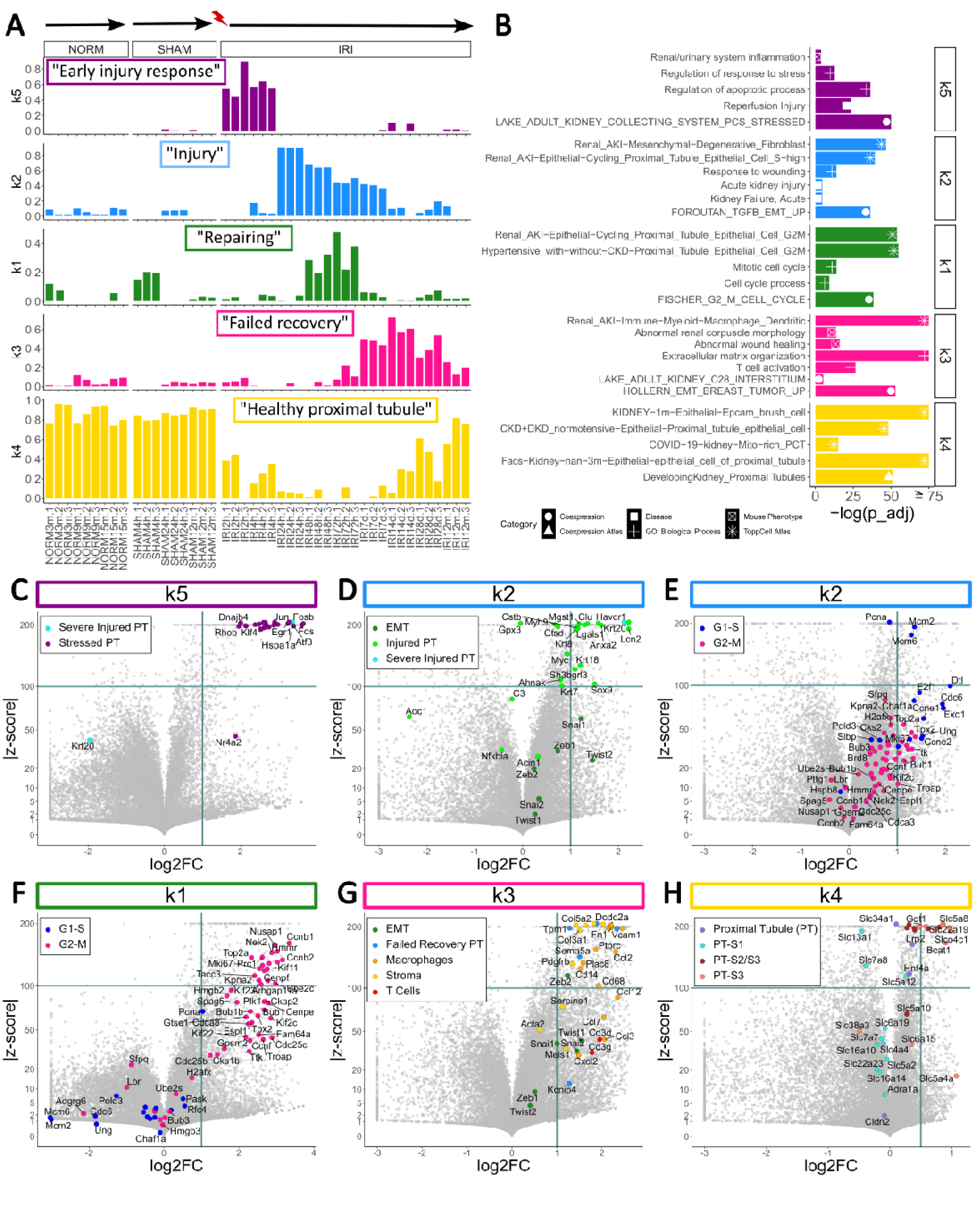
Topic modeling of gene expression levels from a mouse kidney following IRI reveals time-dependent cell states that correspond to early injury response, injury, repairing, failed recovery, and healthy proximal tubule. (A) A structure plot showing the proportions of the K=5 time-dependent cell states (topics) following IRI. Also shown are samples from normal and sham surgery controls. The topics were identified using GO enrichment analysis and known marker genes from the literature. (B) Gene Ontology (GO) enrichment analysis for overexpressed genes in each topic. Overexpressed genes were defined to be those with log2FC > 1 and |z_score| > 100, apart from topic k4 for which we chose log2FC > 0.5. (C–H) Volcano plots showing the differential expression of gene sets associated with known cell states within each topic.

Topic “k5” appears immediately after injury (2-4 hours following IRI, Figure 1A) and over-expresses a set of genes related to stress in the proximal tubule (e.g. the genes Atf3, Dnajb4, Fos, Fosb, Jun, and Rhob from the set labeled as “Stressed PT”) (Liu et al. 2017; Shin et al. 2008) and a marker for severe injury (the gene Hspa1a from the set labeled as “Severe Injured PT”) (Kirita et al. 2020; Musiał and Zwolińska 2011) (Fig 1C). GO enrichment analysis (Figure 1B) showed that this topic significantly over-expresses genes related to renal inflammation, apoptosis, reperfusion injury, and stress. Moreover, we found that this topic also over-expresses immediate-early response genes (Tullai et al. 2007; Winkles 1997) (Figure S1 and Table S3), a gene family known to be activated rapidly and transiently in response to external stimuli, subsequently regulating cell proliferation, differentiation, and apoptosis. Hence, we labeled this topic as “Early injury response”.

Topic “k2” that peaks at 24 hours following IRI and slowly decays until 7 days post-injury (Figure 1A) over-expresses a set of genes related to tubular injury (e.g. the genes Clu, Havcr1, Krt18, and Myc, from the set labeled as “Injured PT”) (Kirita et al. 2020; Liu et al. 2017; Rudman-Melnick et al. 2020) and a marker for severe proximal tubular injury (the gene Krt20, from the set labeled as “Severe Injured PT”) (Kirita et al. 2020; Liu et al. 2017) (Figure 1D). We also found upregulation of a set of genes marking the Epithelial to Mesenchymal Transition (e.g. Snai1, Twist2, and Zeb1, from the gene set labeled as “EMT”) (Figure 1D), and genes that are known to be over-expressed during the G1-S phases of the cell cycle (e.g. Ccne1, Cdc6, Mcm2, and Mcm6, from the gene set labeled as “G1-S”) (Dominguez, Tsai, Gomez, et al. 2016; Dominguez, Tsai, Weatheritt, et al. 2016) (Figure 1E and Table S3). GO enrichment analysis (Figure 1B) showed that this topic significantly over-expresses genes related to injured proximal tubular cells at S-phase, response to wounding, acute kidney injury, and EMT. Hence, we labeled this topic as “Injury”.

Topic “k1” is elevated at 48 to 72 hours after injury (Figure 1A) and over-expresses a set of genes known to be involved in the G2-M phases of the cell cycle (e.g. Ccnb1, Ccnb2, Mki67, and Top2a, from the gene set labeled as “G2-M”) (Dominguez, Tsai, Gomez, et al. 2016; Dominguez, Tsai, Weatheritt, et al. 2016) (Figure 1F and Table S3). GO enrichment analysis (Figure 1B) showed that this topic significantly over- expresses genes related to cycling proximal tubular cells at the G2-M phases of the mitotic cell cycle. Notably, this topic shows high expression of Top2a together with low expression of many renal tubular markers, consistent with previous characterization of repairing proximal tubular cells identified by Kirita et al. (Kirita et al. 2020) (that also peak at 2 days post-IRI, Figure S2). Therefore, we labeled this topic as “Repairing”.

Topic “k3” emerges at later times, from 7 days to 12 months post-IRI (Figure 1A), and over-expresses a set of genes marking EMT (e.g. Snai2, Twist1, and Zeb2, from the gene set labeled as “EMT”), markers for kidney stroma (e.g. Col3a1, Col5a2, and Fn1, from the set labeled as “Stroma”), genes marking macrophages (e.g. Ccl2, Cd14, Cd68, and Plac8, from the gene set labeled as “Macrophages”) and T cells (e.g. Cd3g and Cd3d from the set “T Cells”), and markers for failed kidney repair (the genes Dcdc2a, Sema5a, Tpm1, and Vcam1, from the gene set labeled as “Failed Recovery PT”) (Kirita et al. 2020; Muto et al. 2021) (Figure 1G). GO enrichment analysis (Figure 1B) showed that topic “k3” significantly over- expresses genes associated with macrophages, T cell activation, abnormal wound healing, extracellular matrix organization, kidney interstitium, and EMT. Hence, we termed this topic as “Failed recovery”.

The last topic, “k4”, which was predominant in normal and sham surgery samples, exhibited a transient decline following IRI and nearly recovered to baseline levels after 12 months (Figure 1A). This topic over- expresses a set of genes marking the healthy proximal tubule (e.g. Hnf4a, Lrp2, and Slc34a1, from the gene set labeled as “Proximal Tubule”) and specific markers for Segments 2 and 3 of the proximal tubule (e.g. Bcat1, Ggt1, Slc5a8, and Slc22a19, from the gene sets labeled as “PT-S2/S3” and “PT-S3”) (Figure 1H). Interestingly, this topic did not over-express markers specific to Segment 1 of the proximal tubule (the gene set labeled as “PT-S1”). GO enrichment analysis (Figure 1B) showed that this topic significantly over-expresses genes that are known to be elevated in the epithelial cells of the kidney proximal tubule. Consequently, we labeled this topic as “Healthy proximal tubule”.

In order to inspect the relation between cell division and repair following kidney injury, we inspected expression levels of the BIRC proteins family. This family consists of the genes BIRC1 (NAIP), BIRC2, and BIRC3, that are involved in inhibition of apoptosis, and BIRC5 which is a regulator of mitotic cell division (Silke and Vaux 2001). We found that Birc5 is highly over-expressed during 48 to 72 hours following injury (Figure S3), similar to the behavior of topic “k1” (that we labeled as “Repairing” and that over- expresses the gene Top2a). Birc2 and Birc3 however, were elevated from 4 hours to 28 days following IRI, and the BIRC1 mouse orthologue, Naip2, was elevated from 48 hours to 12 months following IRI, thus covering the whole duration of repair.

To test the robustness of our results with respect to the selected number of topics, we also fitted a topic model with k=6 topics (on the normalized counts matrix) and obtained similar results (Supplementary information, Figures S4-S5).

RNA transcripts undergo alternative splicing following kidney injury

We next used rMATS (Shen et al. 2014) to identify transcripts that are alternatively spliced following IRI (Figures 2-3 and Figures S7-S23). To this end, we chose time points that represent each of the cell states identified by topic modeling and performed comparisons between them (Figure S7). We identified 85 cassette exons (i.e. skipped exons/SE) that are significantly alternatively spliced (FDR = 0 and |Δ1| > 0.2) in at least one of these comparisons (Figure 2 and Table S9). Then, we manually validated selected transcripts by inspection in the IGV genome browser (Robinson et al. 2011).

**Figure 2:**
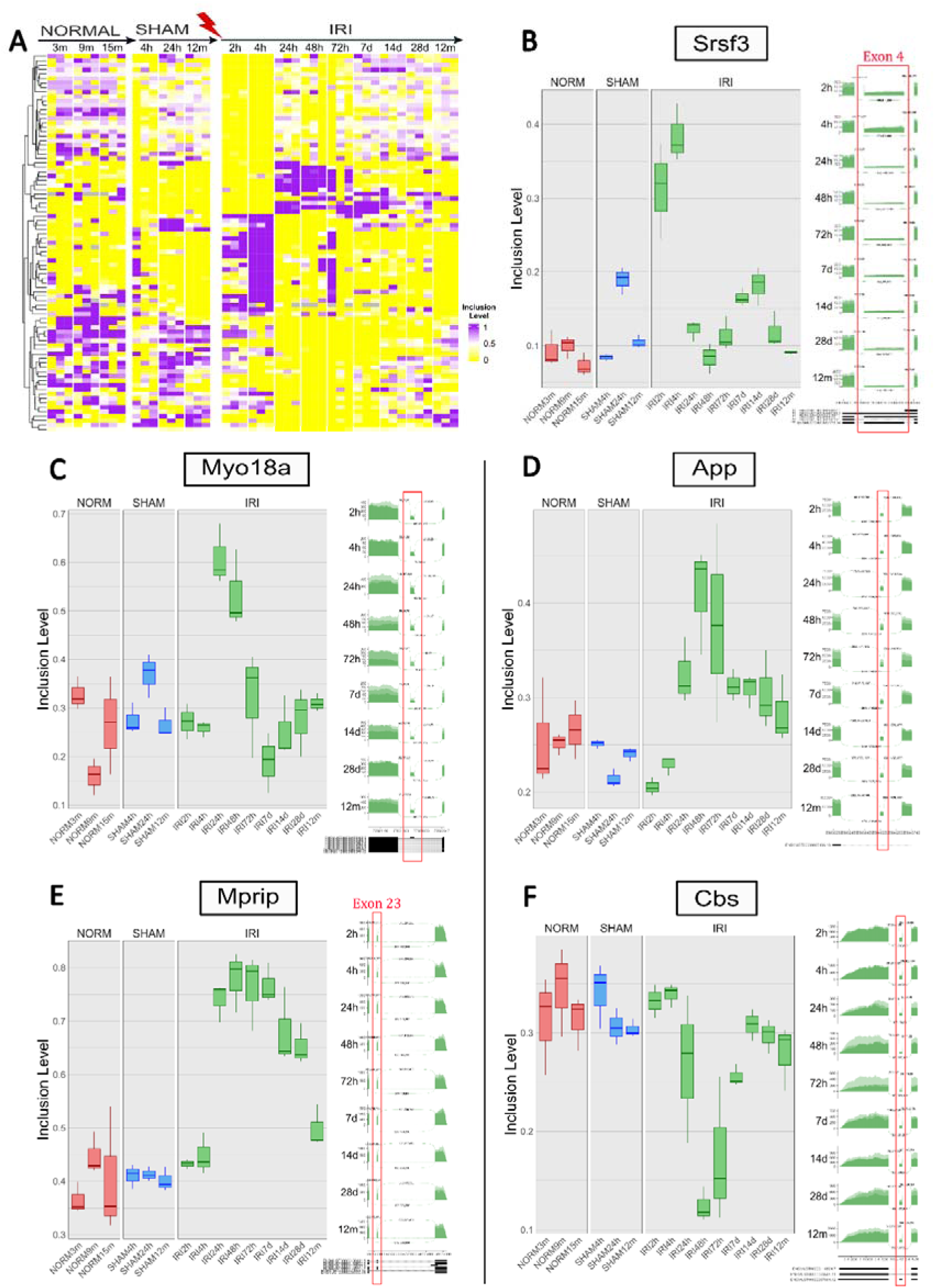
RNA transcripts undergo alternative splicing (exon kipping/inclusion) following kidney injury. (A) A heatmap of exon inclusion levels for 85 cassette exons that were significantly skipped/included in the kidney following IRI, as well as normal and sham surgery controls. We selected exons for which FDR = 0 and |Δ1| > 0.2 in at least one comparison between pairs of selected time-points. (B–F) Boxplots of inclusion levels and sashimi plots for selected cassette exons. Each time-point consists of three replicate samples. Note that the Sashimi plots were drawn only for the samples following IRI and not for the normal and sham surgery controls.

**Figure 3:**
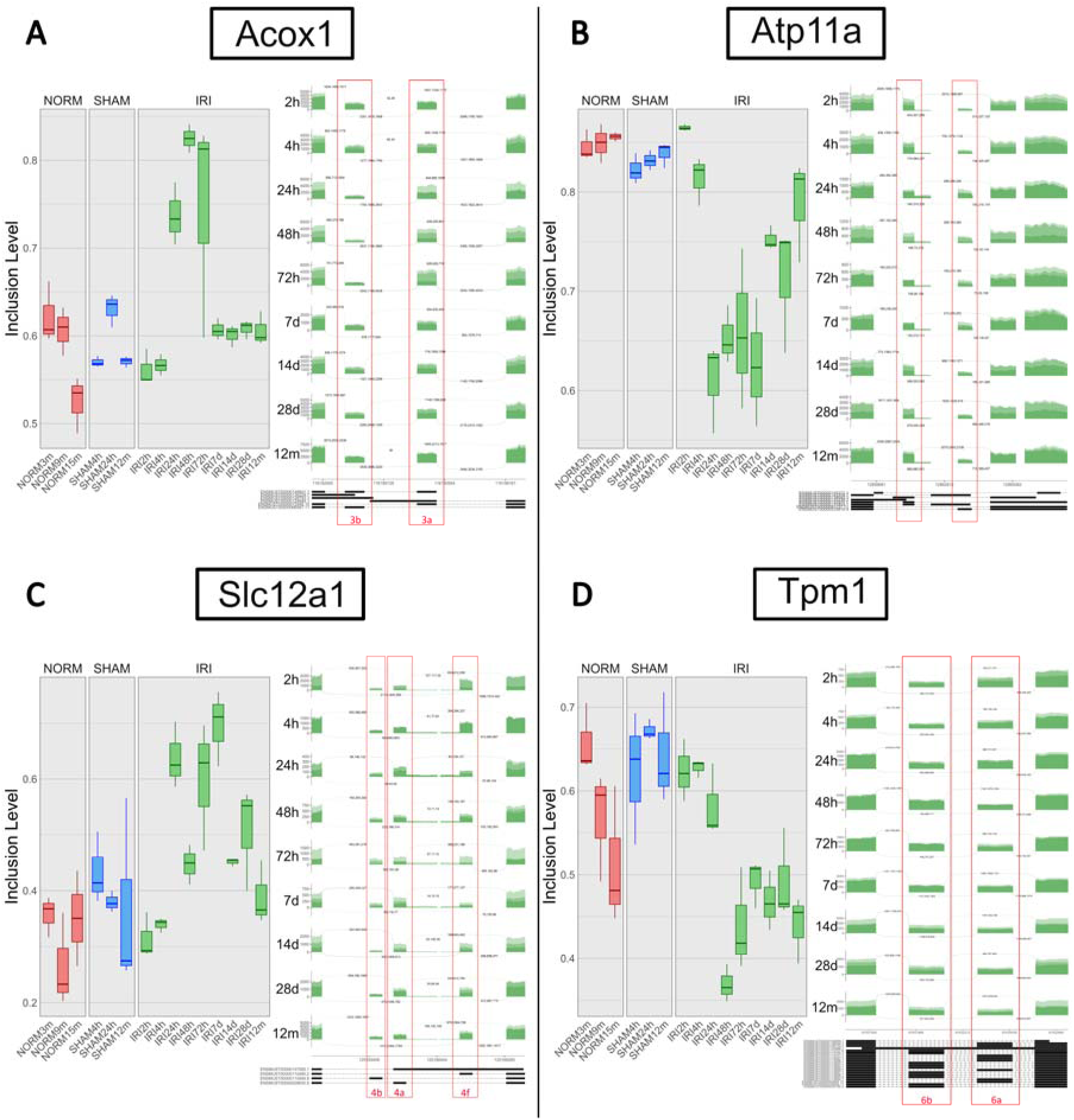
RNA transcripts undergo alternative splicing (mutually exclusive exons, MXE) following kidney injury. (A-D) Boxplots of inclusion levels and sashimi plots for selected mutually exclusive exons that are alternatively spliced following IRI. We selected MXEs for which FDR = 0 and |Δ1| > 0.1 in at least one comparison between pairs of selected time points. Each time-point consists of three replicate samples. Note that the Sashimi plots were drawn only for the samples following IRI and not for the normal and sham surgery controls.

Some of these exons were previously found to be related to injury, stress, and EMT. For example, in the dataset that we studied, inclusion levels of exon 4 of the gene Srsf3 are highly upregulated at 2 and 4 hours post-IRI (Figure 2B, Figure S8). This exon is thought to serve as an autoregulatory mechanism (J. Guo, Jia, and Jia 2015; Jumaa 1997) since it contains a premature stop codon, whose inclusion is promoted by Srsf3 expression. Thus, excessive expression of Srsf3 results in exon 4-including isoforms that are translated into truncated Srsf3 proteins. It was also recently suggested that truncated Srsf3 acts as a positive regulator of oxidative stress-initiated inflammatory response in a colon cancer cell line (Kano et al. 2014).

We also observed a cassette exon of the gene Myo18a with elevated inclusion levels at 24 and 48 hours following injury (Figure 2C). Myo18a has been associated with epithelial cell migration and cell survival following DNA damage (Farber-Katz et al. 2014; Hsu et al. 2010), and also with EMT in breast cancer (Naro et al. 2021) (see Figures S9-S11). Similarly, inclusion levels of exon 11 of the gene Cttn are elevated from 24 hours to 28 days following injury (Figure S12-13). In a recent study, it was found that exon 11 of Cttn is upregulated in a mouse lung adenocarcinoma cell line that undergoes EMT-associated changes in cell morphology and gene expression following stimulation with the EMT regulator TGF-beta (Miyashita et al. 2021). Similar results were found by Braeutigam et al. (Braeutigam et al. 2014), who also identified the RNA binding protein Rbfox2 to be a regulator of this exon. These findings are also consistent with an earlier report that overexpression of CTTN variants lacking exon 11 in cell lines results in reduced cell migration (Van Rossum et al. 2003).

Exon 23 of the gene Mprip shows higher inclusion levels from 24 hours to 28 days following IRI (Figure 2E and Figure S14). It was previously found that inclusion of this exon is induced by the RNA binding protein RBFOX2 in metastatic pancreatic cancer cells (Jbara et al. 2023). Moreover, in the mouse fetal kidney, inclusion of exon 23 was found to be higher in mesenchymal cell populations compared to epithelial cell populations (Wineberg et al. 2020).

We performed a similar analysis for mutually exclusive exons (MXE) that undergo alternative splicing following IRI. Again, we selected MXEs that were significantly alternatively spliced (FDR = 0 and |Δ1| > 0.1) in at least one of the comparisons, and manually validated them by inspection in the genome browser. For example, we observed a significant switching between exons 3a and 3b of the gene Acox1 (Figure 3A and Figure S20). These exons discern between two known splice variants, Acox1a and Acox1b, that include either exon 3a or exon 3b respectively, and are thought to have different substrate specificities (Oaxaca-Castillo et al. 2007; Varanasi et al. 1994). It was found in the kidney that Acox1a mRNA levels are generally higher relative to Acox1b (Vluggens et al. 2010). Here we observed that, following IRI, there is a transient increase of Acox1a relative to Acox1b from 24 to 72 hours following kidney injury, even more than in the normal kidney.

Likewise, we observed a significant switching between exons 4a and 4f of the gene Slc12a1 following IRI (Figure 3C and Figure S21). The gene Slc12a1 encodes the Na-K-2Cl (NKCC2) co-transporter in the epithelial cells of the thick ascending loop of Henle (TAL). At least three of its transcripts differ in their variable exon 4: isoform NKCC2A includes exon 4a, isoform NKCC2B includes exon 4b, and isoform NKCC2F includes exon 4f. The different isoforms were previously found to be expressed in different locations of the thick ascending loop of Henle, where NKCC2A is present both in the medulla and the cortex, NKCC2B is predominantly expressed in the cortex, and NKCC2F is predominantly localized to the medulla (Castrop and Schnermann 2008). It was previously shown that levels of the transcript isoform NKCC2A in the medullary thick ascending loop of Henle increase following hypertonic NaCl intake, and that this, along with expression levels of the gene NFAT5, induces production of TNF (Hao, Bellner, and Ferreri 2013) which is commonly linked to inflammation. Here we observe a significant switching between NKCC2A and NKCC2F, with a higher proportion of NKCC2A relative to NKCC2F from 24 hours to 7 days following IRI.

We also found that the mutually exclusive exons 6a and 6b of the gene Tpm1 undergo switching following IRI (Cao, Routh, and Kuyumcu-Martinez 2021) (Figure 3D and Figure S22). In a previous study we found that the inclusion of exon 6b relative to exon 6a was associated with the more mesenchymal populations of the developing fetal kidney, i.e. the nephrogenic zone stroma (un-induced mesenchyme) and the cap mesenchyme (Wineberg et al. 2020). In the present study we observed that exon 6b is significantly elevated relative to exon 6a from 48 hours until 12 months following IRI, indicating the long-term persistence of an injury-related stromal component. We also found an additional splicing variant in the 5’ end of the transcript (Figure S23), where we observed that exons 1a and 2b (which are associated with transcripts “Tpm1.6/7” (Xu et al. 2024)) are upregulated relative to exon 1b (which is associated with transcripts “Tpm1.8/9”) from 48 hours to 14 days following IRI. This upregulation was shown to be associated with increased cell proliferation (Xu et al. 2024) in ovarian cancer cell lines. Note that TPM1 was previously observed to be over-expressed in the “PT_VCAM1” cell population, a cell population over-expressing the gene VCAM1 that is thought to be associated with proximal tubular cells that fail to repair following kidney injury (Muto et al. 2021).

Finally, we performed Gene Ontology (GO) enrichment analysis for genes whose cassette exons were found to be significantly alternatively spliced between different time points (FDR = 0 and |Δ1| > 0.2), and found that many of these transcripts are related to the regulation of apoptosis (Table S9). This is consistent with upregulation of the apoptosis inhibitors Birc1 (Naip), Birc2, and Birc3 up to 12 months following kidney injury (Figure S3).

Motif enrichment analysis for RNA binding proteins (RBPs) uncovers putative splicing regulators

We used rMAPS to identify putative splicing regulators (Hwang et al. 2020; Park et al. 2016). rMAPS accepts a list of alternatively spliced exons as input and searches their surrounding genomic regions for RNA binding motifs associated with known splicing regulators (Figure 4, Table S10, and Figures S24-S30). This yielded a set of putative splicing regulators whose binding motifs were enriched upstream or downstream of exons that were alternatively spliced, and that were also differentially expressed between different time points following IRI.

**Figure 4:**
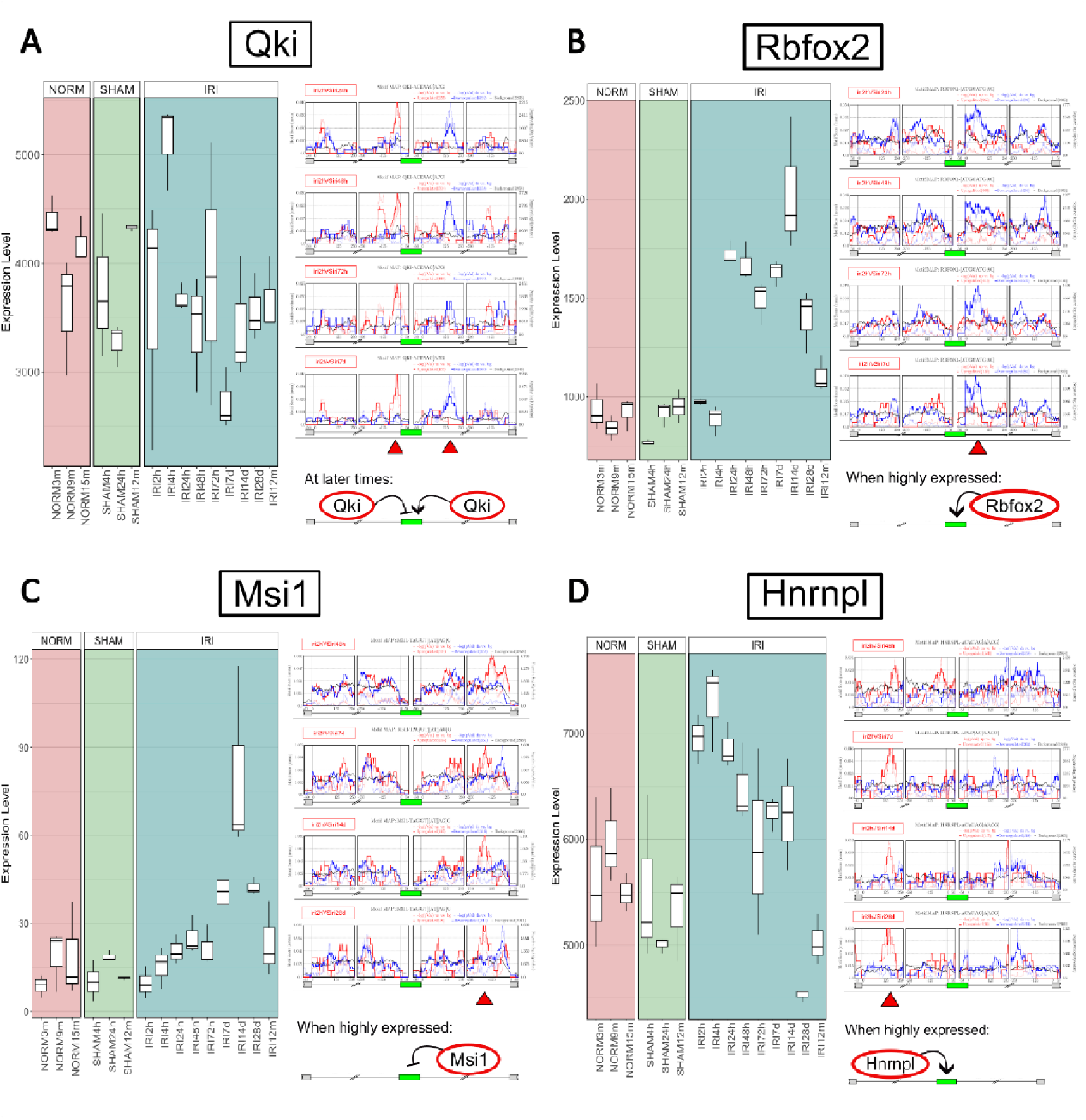
Motif enrichment analysis for RNA binding proteins reveals putative splicing regulators. Shown are motif enrichment diagrams (right) and normalized gene expression boxplots (left) for selected splicing regulators. (A) Motif enrichment analysis indicates that at later times following IRI, the RNA binding protein Qki suppresses exon inclusion by binding to upstream introns and promotes exon inclusion by binding downstream. (B) Motif enrichment analysis indicates that Rbfox2, which is a known regulator of EMT-associated splicing, promotes exon inclusion when it is over-expressed, by binding to downstream introns. (C) Motif enrichment analysis indicates that Msi1, when over-expressed, promotes exon skipping by binding to downstream introns. (D) Motif enrichment analysis indicates that HNRNPL, when over-expressed at early time points following IRI, promotes exon inclusion by binding to upstream introns.

For example, binding motif enrichment for the splicing regulator Qki (Figure 4A) indicates that at later times following injury, Qki suppresses inclusion of cassette exons by binding to their upstream introns, and promotes their inclusion by binding downstream, concurrent with previous findings of Qki- controlled splicing events associated with EMT (Pillman et al. 2018). Likewise, motif enrichment for the RNA binding protein Rbfox2 (Figure 4B) indicates that, when over-expressed, Rbfox2 promotes exon inclusion by binding to the downstream intronic regions (Yeo et al. 2009). We note that the over- expression of Rbfox2 matches the increased inclusion levels of exon 11 of the gene Cttn (Figure S12), whose inclusion was previously found to be promoted by Rbfox2 and to be associated with EMT (Braeutigam et al. 2014), and also to the increased inclusion levels of exon 23 in the gene Mprip (Figure 2E), whose inclusion was also previously found to be promoted by Rbfox2 (Jbara et al. 2023). Similarly, binding motif enrichment and elevated expression of Msi1 (Figure 4C), a splicing regulator that was previously found to be highly expressed in kidney fibrosis (Jadhav et al. 2016), indicate that, when over- expressed, Msi1 promotes exon skipping by binding downstream of target exons.

## DISCUSSION

In this study we applied topic modeling, an un-supervised machine learning technique, to identify dynamic cell states following kidney injury. Additionally, we identified sets of transcripts that undergo alternative splicing between these cell states, along with their putative splicing regulators. Understanding mechanisms of mRNA splicing may assist in the development of RNA-based technologies for detecting, treating, and monitoring complex diseases (Baughn et al. 2023; Kim et al. 2023). We believe that this study could pave the way for new molecular approaches to monitor and treat kidney injury.

In recent years single-cell RNA sequencing technologies have enabled deep and precise characterization of cell states. However, most of the current high throughput single-cell RNAseq protocols are based on reverse transcription with a poly-T primer containing a cell barcode and a UMI, followed by template switching, cDNA amplification, fragmentation, and short-read sequencing. In order to cover the cell barcode and UMI, only fragments that include the 3’ end of the original transcript are sequenced. As a result, there is typically a strong coverage bias to the 3’ end and insufficient coverage of exon junctions along the whole transcript, which makes it difficult to perform statistically reliable mRNA splicing analysis. While “bulk” RNA sequencing provides much higher coverage along the whole transcript, these measurements cannot discern between the heterogeneous cell populations within the injured kidney, and thus cannot resolve the various cellular and molecular mechanisms that are activated following IRI. To mitigate these shortcomings, we first used topic modeling to “deconvolve” the major cell states within time-dependent “bulk” RNA sequencing measurements, and then used rMATS to identify transcripts that are significantly alternatively spliced between bulk RNA measurements from time points that most represent these major cell states. We believe that our study will provide a stepping stone for future single-cell studies aimed at understanding mechanisms of alternative mRNA splicing in kidney injury and repair.

Recently, technologies for single-cell long-read RNA sequencing were developed by Pacific Bioscience (PacBio) (Al’Khafaji et al. 2023) and Oxford Nanopore (ONT) (Byrne et al. 2017; Lebrigand et al. 2020).

Although these technologies can resolve alternatively spliced transcripts, they still suffer from technical difficulties, such as relatively low read throughput, limiting the number of cells analyzed. This is especially important for analyzing kidney injury and repair, which involves multiple cell populations of varying sizes, some of which are very small. Another difficulty is that, due to the fragility of injured cells, tissue dissociation into a single-cell suspension carries a risk of significant cell loss and contamination of single-cell transcripts from ambient RNA.

It has been observed that several of the genes that are over-expressed or alternatively spliced following kidney injury are related to EMT (e.g. Twist (Kida et al. 2007), see Figures 1D and 1G). We note that although it has been hypothesized that injured kidney tubular cells undergo an EMT-like transition into mesenchymal cells which participate in the development of fibrosis, conclusive in-vivo evidence is still lacking, and the mechanisms involved are yet unclear (Kriz, Kaissling, and Le Hir 2011; Sheng and Zhuang 2020). In the bulk RNA sequencing dataset that we studied we could not observe a clear alternative splicing in the genes Cd44 (Brown et al. 2011; Sneath and Mangham 1998), Ctnnd1 (Warzecha, Sato, et al. 2009; Warzecha, Shen, et al. 2009), Enah (Di Modugno et al. 2012; Shapiro et al. 2011; Warzecha, Shen, et al. 2009), and Fgfr2 (Hovhannisyan, Warzecha, and Carstens 2006; Warzecha, Sato, et al. 2009) (Figures S31-S34), which are frequently alternatively spliced during EMT. However, a better resolution in detecting and quantifying these and similar mRNA splicing isoforms might be obtained by long-read scRNA-seq.

## METHODS

### RNA sequencing data and preprocessing

A total of 49 fastq files were downloaded from the Gene Expression Omnibus (accession number GSE98622) using the SRA toolkit (Table S1). The sequences were aligned to the mouse genome (mm10) using the STAR aligner (Dobin et al. 2013) to obtain the raw counts matrix. To avoid technical bias, we removed 4 samples that were collected 6 months following IRI (SRR5515120, SRR5515121, SRR5515122, and SRR5515123) since they were sequenced on a different machine and with different read lengths from the other samples in the IRI time series. Normalized counts for the remaining 45 samples were obtained using DESeq2 (Love, Huber, and Anders 2014). Genes with zero counts across all samples were removed from the analysis.

### Topic modeling

Topic modeling is an unsupervised machine learning technique that can be used to identify hidden components, or “topics”, in complex datasets (Dey, Hsiao, and Stephens 2017; Girdhar, Giguère, and Dudek 2013; Varadarajan and Odobez 2009). Given a set of documents that are assumed to be composed of k latent topics, the aim of this technique is to discover these k topics (that is, the probability of occurrence of each word in each topic) as well as their proportions in each document (Blei, Ng, and Jordan 2003). In our case, each time-dependent RNAseq expression profile (“document”) is assumed to be composed of a weighted sum of k latent cell states (“topics”). Therefore, by fitting a topic model we can identify the k underlying cell states (that is, the probability of expression of each gene in each topic) as well as their proportions in each RNAseq expression profile.

We performed topic modeling using the function fit_topic_model() from the “fastTopics” R package (Carbonetto et al. 2021; Dey, Hsiao, and Stephens 2017). The outputs of this function are: (1) The “Loadings” L matrix which contains the proportions of the k topics p(topic_k|sample) in each of the samples; (2) The “Factors” F matrix which contains the probability p(gene|topic_k) of a transcript from a given gene being expressed in each one of the k topics (Table S2). Since it is difficult to know the “real” or “optimal” number of topics k a-priori, we first performed topic modeling with k=5 topics on the raw counts matrix. We then repeated our analysis with k=6 topics on the DESeq2-normalized counts and obtained similar results.

To infer the biological identity of the different topics, we checked the expression levels of selected genes that are known from the literature to mark the different cell populations or cell states in the healthy and injured kidney (Table S3). We further characterized the identity of each topic by performing gene ontology (GO) enrichment analysis using ToppGene (Chen et al. 2007) for overexpressed genes. For k=5, overexpressed genes in each topic were defined to be those with log2FC > 1 and |z_score| > 100, apart from topic k4 for which we chose log2FC > 0.5. The log-fold changes between topics were computed using the de_analysis() function with the log-fold change statistics parameter set to lfc.stat = “vsnull”.

Alternative splicing and RNA binding motif enrichment analysis

We used rMATS (Shen et al. 2014) to find genes that are alternatively spliced with significant inclusion level differences between different time-points. For the comparisons, we chose time points that represent the different cell states identified by topic modeling. The IGV genome browser (Robinson et al. 2011) and the ggsashimi command-line tool (Garrido-Martín et al. 2018) were used to visualize and manually inspect alternatively spliced transcripts. To obtain the inclusion levels of all splicing events for all time points in the dataset, we reran rMATS for all samples with the command-line option “–cstat 0”. GO enrichment analysis for significantly alternatively spliced genes was done using ToppGene (Chen et al. 2007) and the Gene Ontology Resource (Ashburner et al. 2000; The Gene Ontology Consortium et al. 2023; Thomas et al. 2022).

In order to identify putative splicing regulators, we used the rMATS output tables as input to rMAPS (Hwang et al. 2020; Park et al. 2016). rMAPS searches for binding motifs belonging to known RNA binding proteins (RBPs) and inspects if they are enriched near the alternatively spliced exons detected by rMATS. The list of RNA binding proteins was obtained from the rMAPS website (http://rmaps.cecsresearch.org/Help/RNABindingProtein) (Table S10). We also followed Yang et al. (Yang et al. 2016) and the CISBP-RNA database (Ray et al. 2013) (http://cisbp-rna.ccbr.utoronto.ca) and assumed that the proteins RBFOX1 and RBFOX2 both bind to the same mRNA motif ([AT]GCATG[AC]).

## ACKNOWLEDGMENTS

We wish to thank Benjamin Humphreys, Yishai Yehuda, Yaron Trink, Tom Maimon, Yochai Israelashvili, Zeev Cohen, and all members of our lab for helpful comments and suggestions.

## AUTHOR CONTRIBUTIONS

Study initiation and conception – D.M. and T.K.; Data analysis – D.M. and T.K.; Other intellectual contribution – J.G.; Manuscript writing – D.M. and T.K.

## DECLARATION OF GENERATIVE AI AND AI-ASSISTED TECHNOLOGIES IN THE WRITING PROCESS

During the preparation of this work the authors used ChatGPT (OpenAI) in order to improve readability and language (e.g. refine sentence structure, wording, and grammar). After using this tool, the authors reviewed and edited the content as needed and take full responsibility for the content of the publication.

## APPENDICES

Supplementary information: Supplementary text and figures.

Table S1: Gene expression counts and metadata obtained from the RNAseq dataset generated by Liu et al.

Table S2: The topics discovered by topic modeling (“factors”) and their proportions at each time point (“loadings”).

Table S3: A list of genes compiled from the literature marking cell subtypes and cell states in the healthy and injured kidney.

Table S4: GO enrichment analysis for genes that are over-expressed in topic k1.

Table S5: GO enrichment analysis for genes that are over-expressed in topic k2.

Table S6: GO enrichment analysis for genes that are over-expressed in topic k3.

Table S7: GO enrichment analysis for genes that are over-expressed in topic k4.

Table S8: GO enrichment analysis for genes that are over-expressed in topic k5.

Table S9: A list of cassette exons that were found to be significantly alternatively spliced following IRI (FDR = 0 and |Δ1| > 0.2, Figure 2A), along with their inclusion levels and results from GO enrichment analysis.

Table S10: A list of known RNA binding proteins and motifs that were examined in this study. Program: A compressed directory containing programs and datasets for data visualization.

