## Supplementary text and figures for "Dynamics of cell states and alternative splicing following kidney ischemia-reperfusion injury"

**Topic modeling with k=6 topics reveals cells states that are similar to those identified with k=5 topics:** In order to check the robustness of our results, we fitted a topic model with k=6 topics to the DESeq2-normalized counts. and obtained results that are similar to those obtained with k=5 topics.

Topic "k6" appears immediately after injury 2-4 hours following IRI (Figure S4A) and over-expresses a set of genes related to stress in the proximal tubule (e.g. the genes Atf3, Dnajb4, Fos, Fosb, Jun, and Rhob from the gene set labeled as "Stressed PT" ) (Liu et al. 2017; Shin et al. 2008) and a marker for severe injury (the gene Hspa1a from the set labeled as "Severe injured PT") (Kirita et al. 2020a; Musiał and Zwolińska 2011) (Fig S4C and Table S3). GO enrichment analysis (Figure S4B) shows that this topic significantly over-expresses genes related to reperfusion injury, apoptosis, stress, and inflammation. Hence, we labeled this topic as "Early injury response".

Topic "k1" peaks at 24 hours following IRI and slowly decays until 7 days post injury (Figure S4A). It over-expresses a set of genes related to tubular injury (e.g. the genes Clu, Havcr1, Krt18, and Myc, from the set "Injured PT") (Kirita et al. 2020a; Liu et al. 2017; Rudman-Melnick et al. 2020) and a marker for severe proximal tubular injury (the gene Krt20, from the set labeled as "Severe injured PT") (Kirita et al. 2020a; Liu et al. 2017) (Figure S4D). We also found upregulation of a set of genes marking the kidney stroma (e.g. Col5a2, Serpine1, and Pdgfrb, from the gene set labeled as "Stroma") (Figure S4D), and genes that are known to be over-expressed during the G1-S and G2-M phases of the cell cycle (Dominguez, Tsai, Gomez, et al. 2016; Dominguez, Tsai, Weatheritt, et al. 2016) (Figure S4E). GO enrichment analysis (Figure S4B) showed that this topic significantly over-expresses genes related to AKI-associated degenerative fibroblasts, injured proximal tubular cells at S-phase, epithelial differentiation, acute kidney injury, and abnormal wound healing. Hence, we labeled this topic as "Injury".

Topic "k3" is elevated at 48 to 72 hours after injury (Figure S4A) and over-expresses a set of genes known to be involved in the G2-M phases of the cell cycle (e.g. Ccnb1, Mki67, Nek2, and Top2a, from the gene set labeled as "G2-M") (Dominguez, Tsai, Gomez, et al. 2016; Dominguez, Tsai, Weatheritt, et al. 2016) (Figure S4F). GO enrichment analysis (Figure S4B) showed that this topic significantly over-expresses genes related to cycling proximal tubular cells at the G2-M phases of the mitotic cell cycle.

Since this topic shows high expression of Top2a, we labeled this topic as “Repairing”, consistent with Figure 2A in Kirita et al. (Kirita et al. 2020a).

Topic “k5” emerges at later times, from 7 days to 12 months post IRI (Figure S4A), and over-expresses a set of genes marking macrophages (e.g. C1qa, C1qb, C1qc, and Plac8, from the gene set “Macrophages”) and T cells (e.g. Cd3d and Cd3g from the gene set “T Cells”), genes marking EMT (e.g. Snai2, Twist1, and Zeb2, from the gene set labeled “EMT”), markers for kidney stroma (e.g. Col1a1, Col5a2, and Fn1, from the gene set “Stroma”), and markers for failed kidney repair (the genes Dcdc2a, Sema5a, Tpm1, and Vcam1, from the gene set “Failed Recovery PT”) (Kirita et al. 2020a; Muto et al. 2021) (Figure S4G-H). GO enrichment analysis (Figure S4B) showed that topic “k5” significantly over-expresses genes associated with chronic kidney failure, increased T cell number, EMT, kidney stroma, and macrophages. Hence, we labeled this topic as “Failed recovery”.

Topic “k4”, was found to be dominant only in the normal kidney samples (Figure S4A). This topic over-expresses a set of genes marking segments 2 and 3 of the proximal tubule (e.g. Bcat1, Ggt1, Slc7a13, and Slc9a8, from the gene sets “PT-S2/S3” and “PT-S3”) (Figure S4I). Interestingly, this topic did not over-express markers specific to segment 1 of the proximal tubule (gene set “PT-S1”). We labeled this topic as “Healthy proximal tubule 1”. Another topic, “k2”, was found to be dominant in both normal and sham surgery samples, to transiently decay following IRI, and to recover almost completely after 12 months (Figure S4A). In contrast to topic “k4”, topic “k2” over-expresses genes marking all three segments of the proximal tubule (sets “Proximal Tubule (PT)”, “PT-S1”, ”PT-S2/S3”, and “PT-S3”) (Figure S4J). GO enrichment analysis (Figure S4B) showed that both topics “k4” and “k2” significantly over-express genes that are known to be elevated in the kidney proximal tubular epithelium, as well as in adjacent epithelial compartments such as podocytes and the loop of Henle. The difference between topics “k2” and “k4” does not seem to originate from the presence of other kidney cell sub-populations (such as the loop of Henle, Figure S5). It is possible that topic “k4” represents an experimental bias unique to the normal samples, due to the fact that they were sequenced on a different sequencing machine and with a different read length compared to the other samples (Table S1). Note also that the DESeq size factors differ in the normal samples and have a pattern that is somewhat similar to topic “k4” (Figure S6).

### SUPPLEMENTARY FIGURES

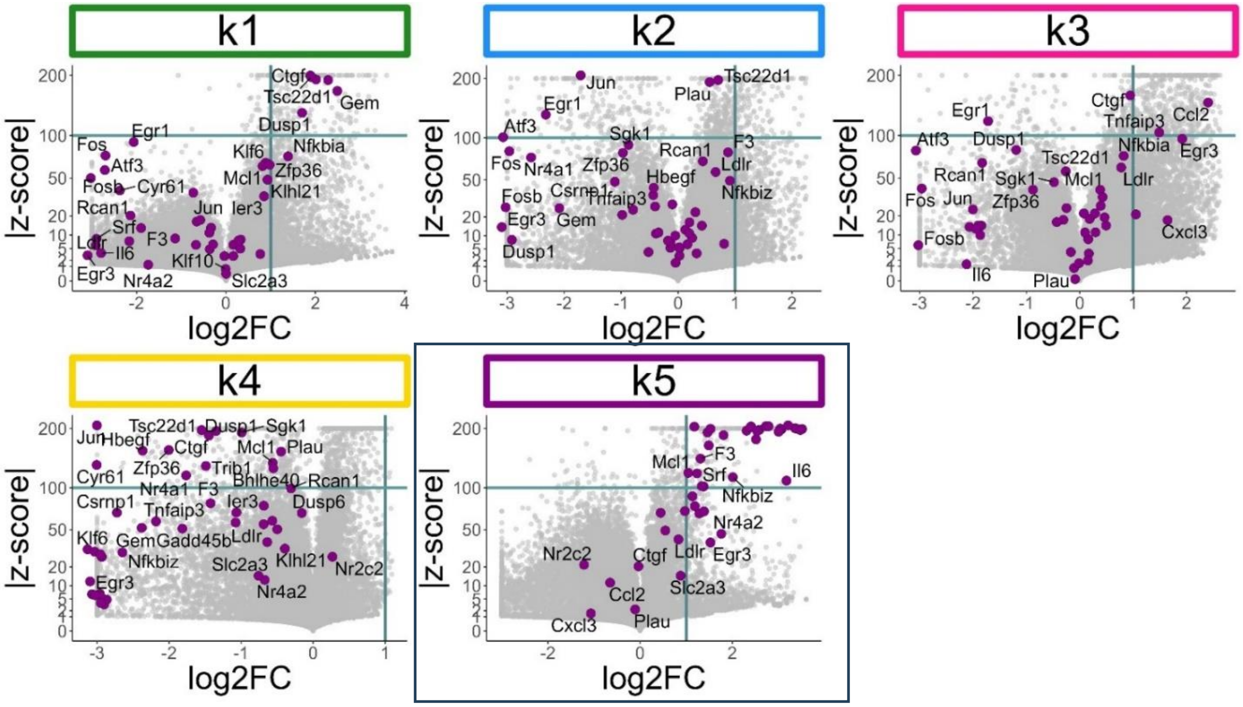


**Figure S1: Most of the Immediate-early response genes are over-expressed in topic k5 (k=5 topics) that we identified as "Early injury response".**

Shown are volcano plots for the different topics, where the Immediate-early response genes as identified by Tullai et al. are marked (Tullai et al. 2007) (Table S3). It can be seen that most of the Immediate-early response genes are over-expressed in topic k5 ("Early injury response", k=5 topics) and under-expressed in topic k4 ("Healthy proximal tubule", k=5 topics).

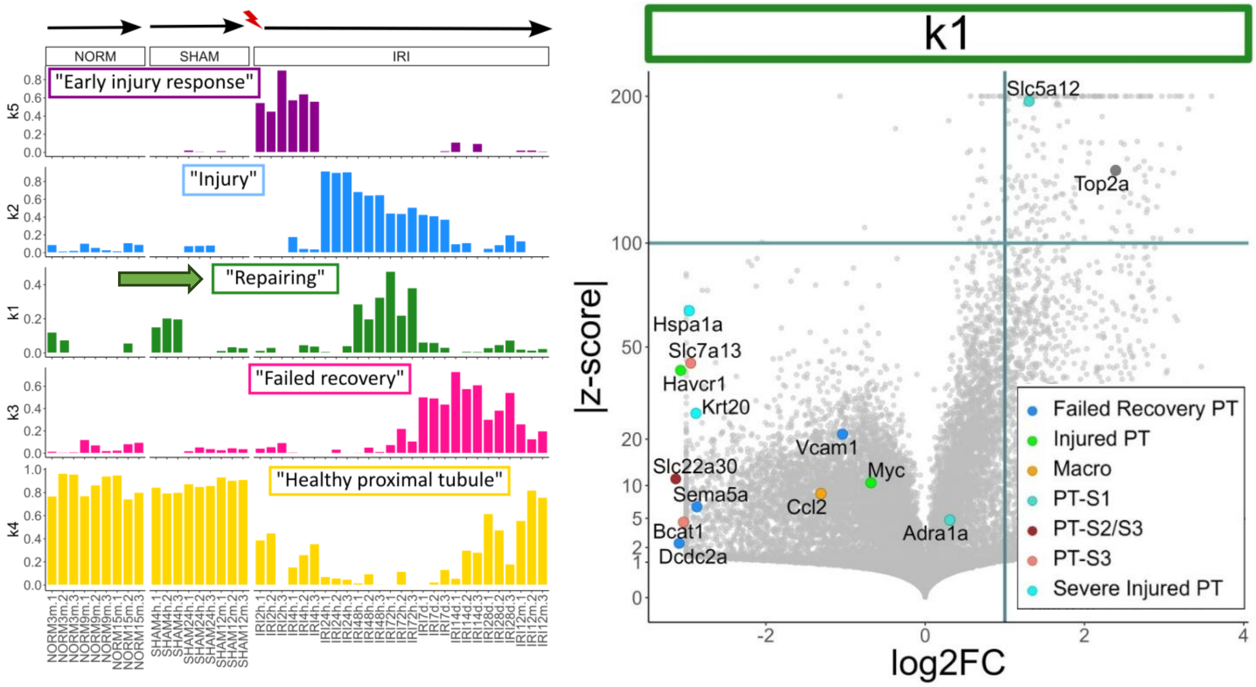


**Figure S2: Topic k1 ("Repairing", k=5 topics) that appears 2 days post IRI overexpresses genes such as Top2a in a way that is similar to the "Repairing PT" cell population identified in Figure 2A of Kirita et al. (Kirita et al. 2020b)**

Shown is a volcano plot of genes over-expressed in topic k1. We marked the genes that appear in the dotplot in Figure 2A of Kirita et al. (Kirita et al. 2020b). These genes mark the cell subpopulations identified therein ("Healthy S1", "Healthy S2", Healthy S3", "Repairing PT", "Injured S1/2", "Injured S3", "Severe injured PT", and "Failed repair PT"). It can be seen that both topic k1 and the "Repairing PT" cell population in Kirita et al. over-express Top2a and under-express the healthy proximal tubular S2/3 markers Slc22a30, Slc7a13, Bcat1, the injured proximal tubular markers Myc, Havcr1, Krt20, Hspa1a, and the failed repair markers Vcam1, Dcdc2a, Sema5a, and Ccl2**.** Note though that there is a slight discrepancy in a single gene - the healthy proximal tubular S1 marker Slc5a12 is over-expressed by topic k1 but has low average expression per cell in the "Repairing PT" cell population in Kirita et al.

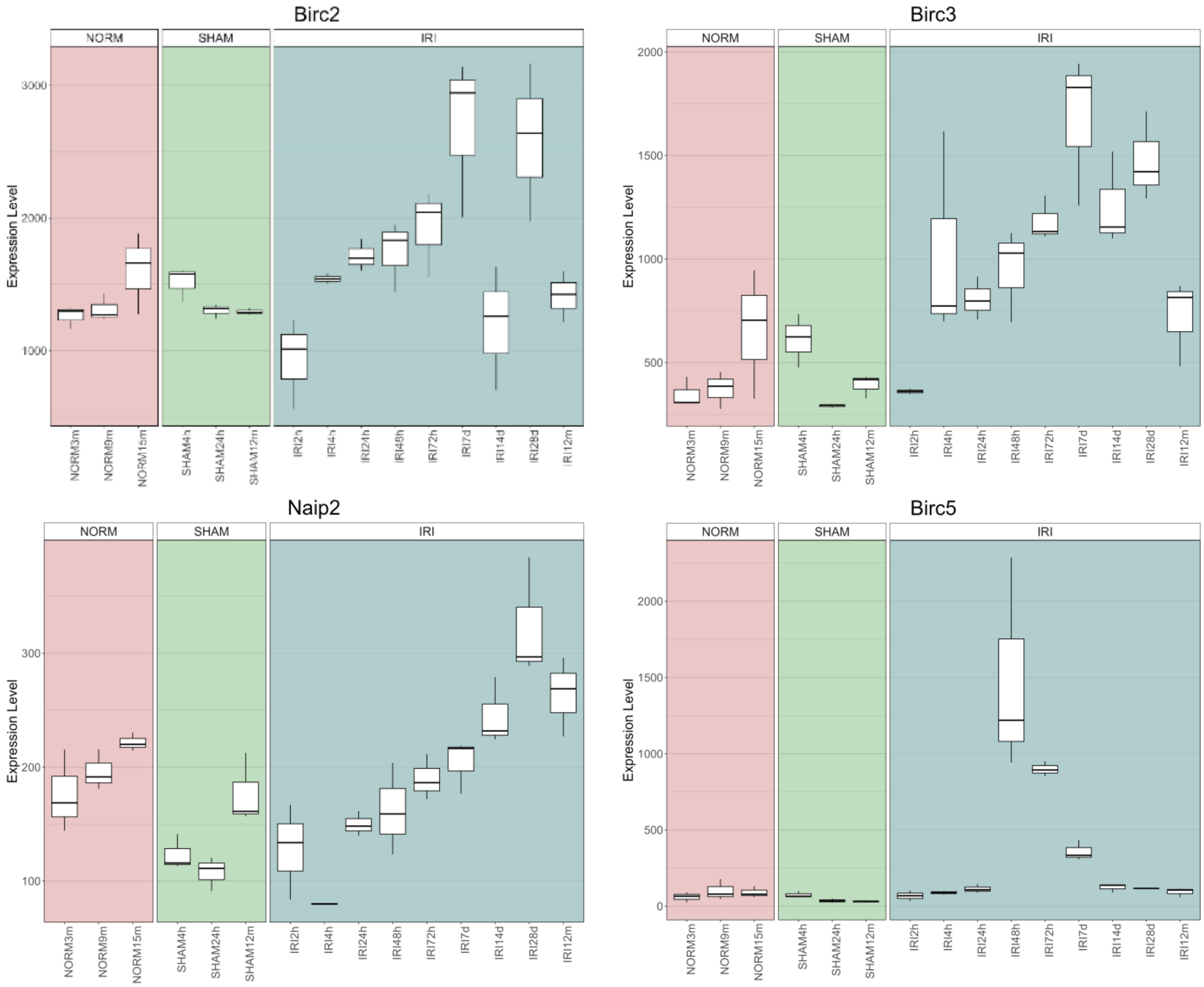


**Figure S3: The anti-apoptotic genes Birc2, Birc3 and Naip2 (Birc1) are elevated throughout the duration of kidney injury response and repair, while the cell cycle related gene Birc5 is over-expressed during 48 to 72 hours following IRI, similar to topic k1 (“Repairing”, k=5 topics).**

Shown are barplots of gene expression levels for the BIRC gene family. It can be seen that Birc2 and Birc3 are elevated from 4 hours to 28 days following IRI and that Naip2 (the mouse ortholog for the human BIRC1/NAIP gene with 76.84% nucleic acid similarity (https://www.genecards.org/cgi-bin/carddisp.pl?gene=NAIP)) is elevated from 24 hours to 12 months, thus covering almost the whole duration of injury response and repair. This is consistent with the presumed anti-apoptotic function of these genes. On the other hand, Birc5, a regulator of mitotic cell division, is highly over-expressed during 48 to 72 hours following injury, similar to the behavior of topic k1 (that we labeled as “Repairing” in the topic model with k=5).

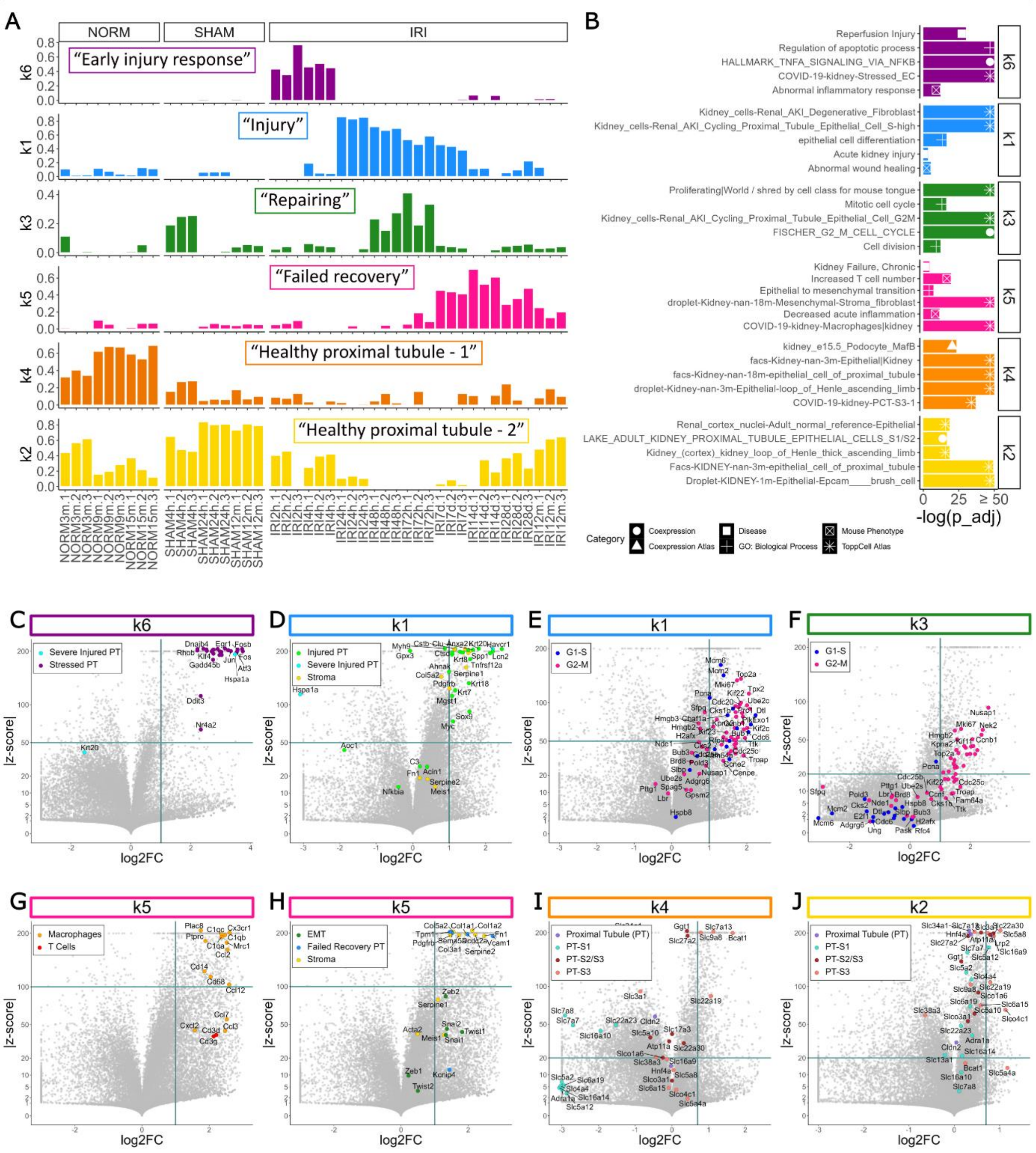


**Figure S4: Topic modeling of gene expression levels from a mouse kidney following IRI with k=6 topics reveals time-dependent cell states that are similar to those identified with k=5 topics.**

(A) A structure plot showing the proportions of k=6 time-dependent cell states (topics) following IRI. Also shown are samples from normal and sham surgery controls. (B) Gene Ontology (GO) enrichment analysis for overexpressed genes in each topic (Overexpressed genes were defined as follows: log2FC > 1 for topics k6, k1, k3, and k5; logFC > 0.7 for topics k4 and k2; |z_score| > 100 for topic k5; |z_score| > 50 for topics k6 and k1; |z_score| > 20 for topics k3, k4, and k2). (C–J) Volcano plots showing the differential expression of gene sets associated with known cell states within each topic.

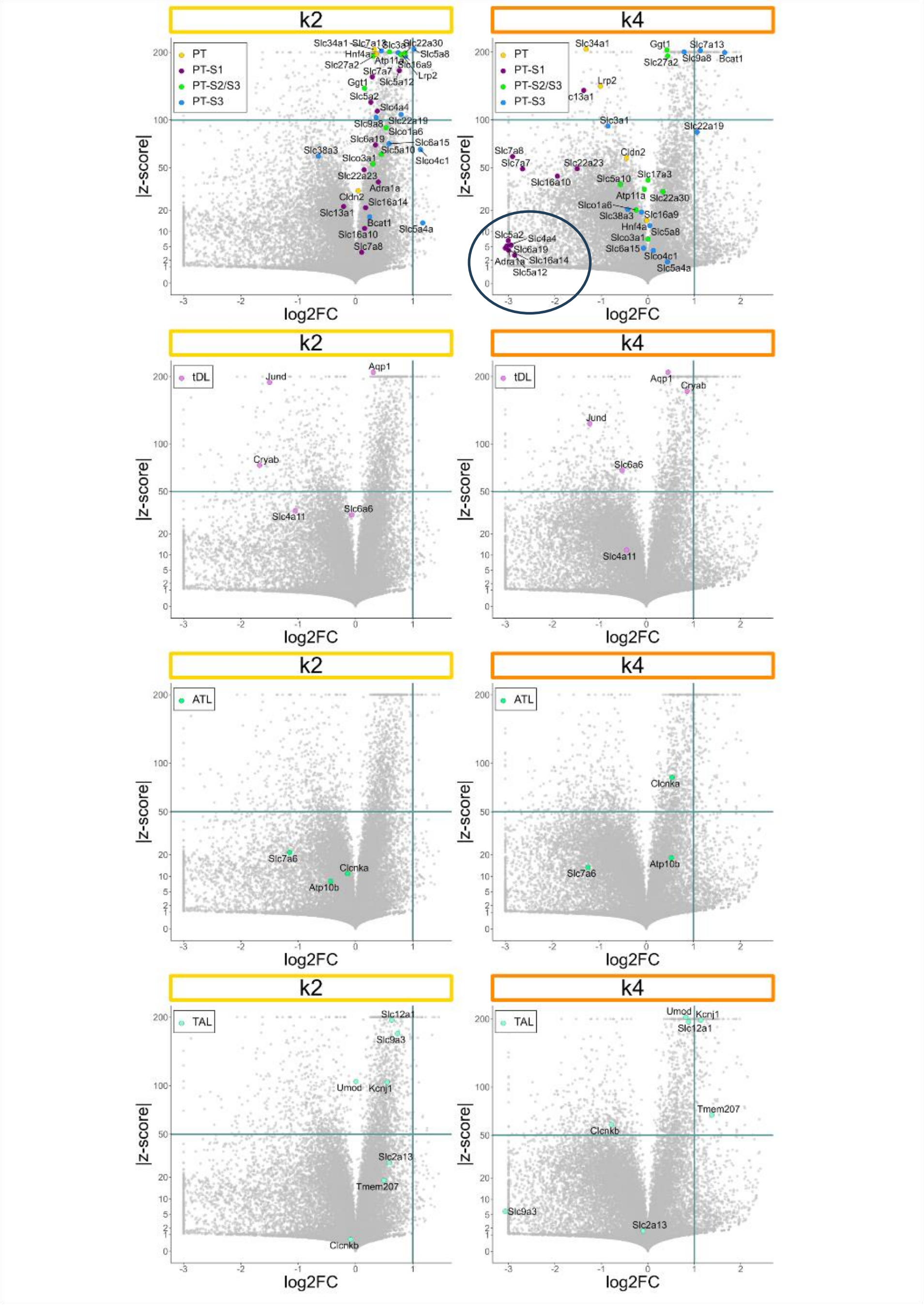


**Figure S5: Topics k2 and k4 for the topic model with k=6 do not show a significant difference in expression for markers of the healthy proximal tubule and the loop of Henle.**

Shown are volcano plots for topics k2 and k4, labeling genes marking the following cell compartments: Proximal Tubule (PT), Proximal Tubule Segment 1 (PT-S1), Proximal Tubules Segments 2 and 3 (PT-S2/S3), Proximal Tubule Segment 3 (PT-S3), thin descending loop (tDL), thin ascending loop (ATL), and the thick ascending loop (TAL). Apart from under-expression of PT-S1 markers in topic k4, we could not discern a significant difference in expression levels of known markers for the proximal tubule and the loop of Henle.

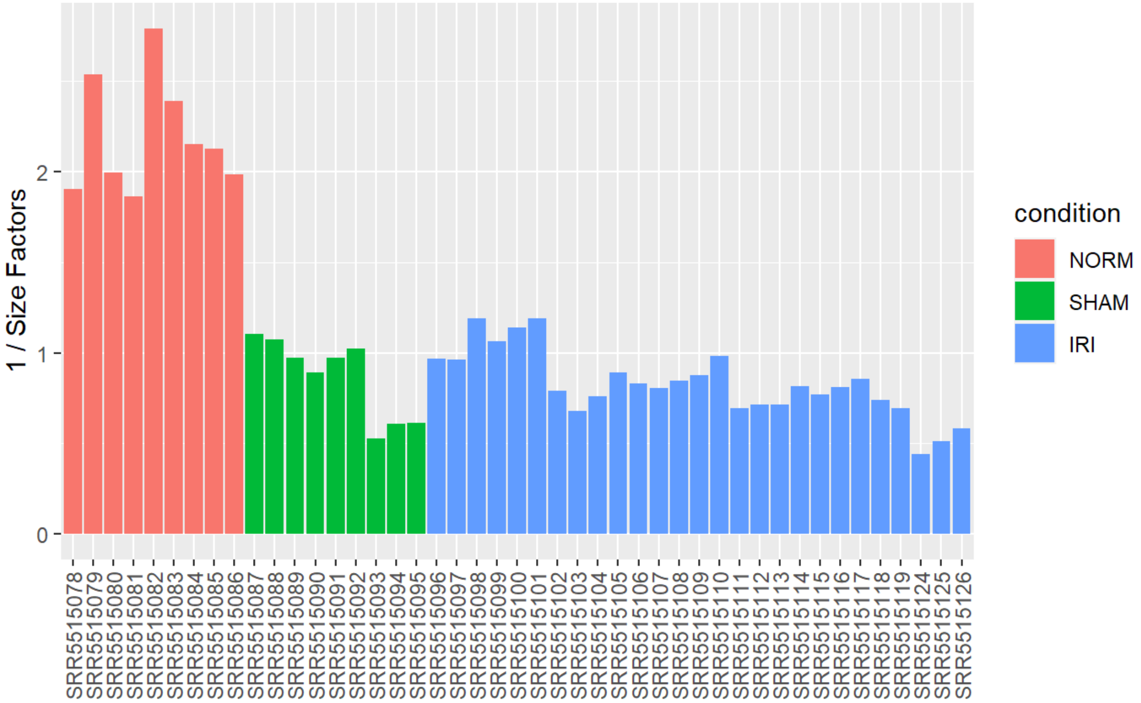


**Figure S6: DESeq2 size factors indicate an experimental bias between the normal samples vs. those following sham surgery and IRI.**

Shown is a barplot of the DESeq2 size factors for all samples. It is seen that size factors for the normal kidney samples are approximately two-fold smaller than the size factors for all other samples. Note also that the sequencing length for the normal samples is shorter (76 bp x 2) than for the samples following sham surgery and IRI (100 bp x 2) and that they were sequenced on a different machine (Liu et al. 2017) (Table S1).

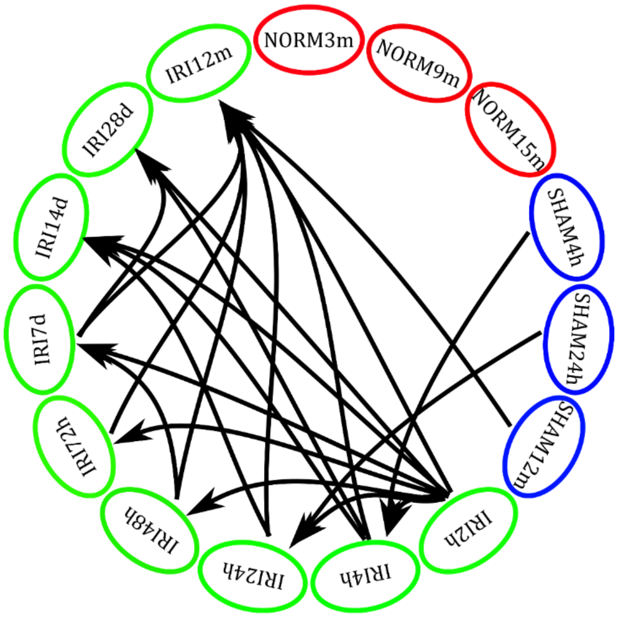


**Figure S7: A sketch of the samples and time-points that were compared by rMATS for identifying alternatively spliced transcripts.**

For the comparisons, we chose time points associated with the different cell states identified by topic modeling.

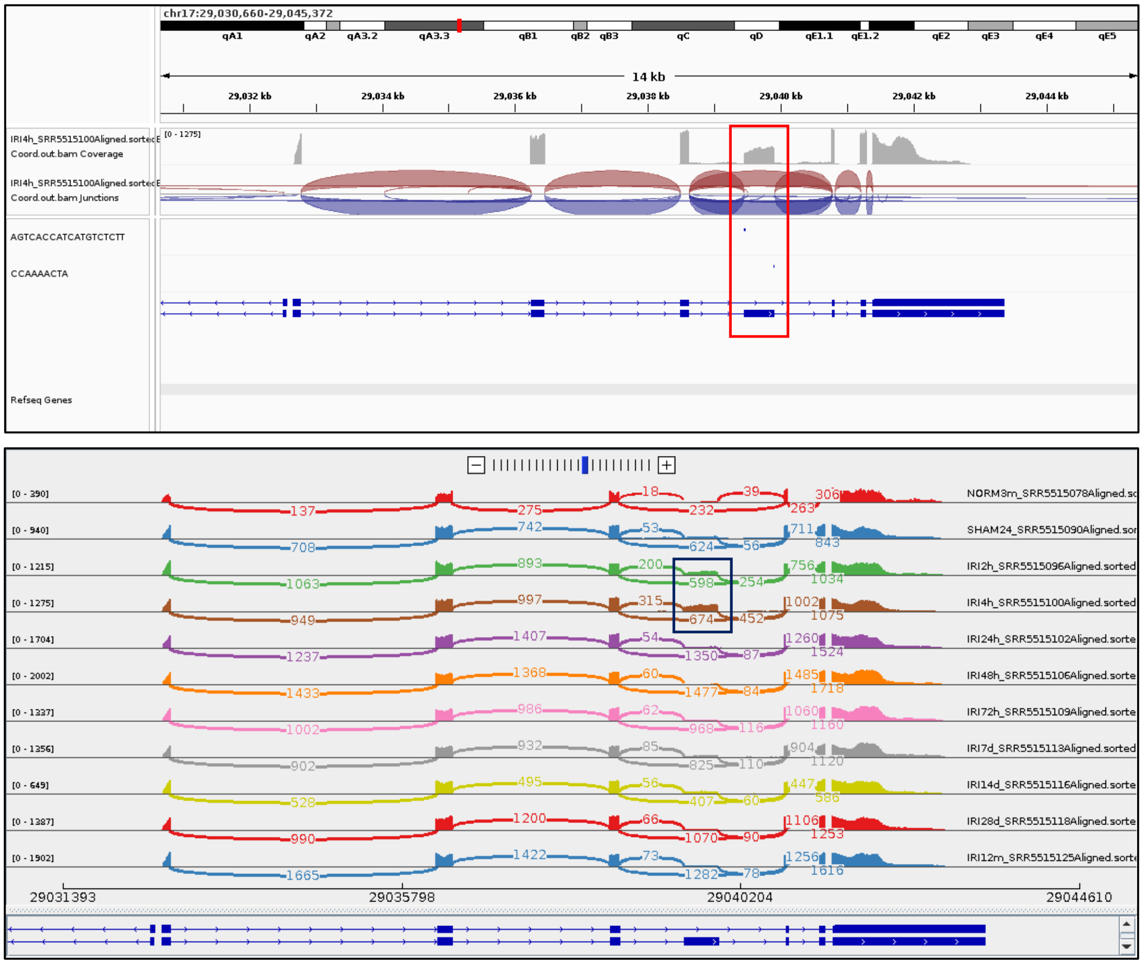


**Figure S8: Exon 4 of the gene Srsf3 is over-expressed at 2 and 4 hours following IRI.**

Top: Identification of the alternatively spliced exon. Shown are 5’ and 3’ subsequences of exon 4 in the mouse Srsf3 gene, as identified in Jumaa et al. (Jumaa 1997). Bottom: A sashimi plot of selected representative samples showing elevated inclusion levels of exon 4 at 2 and 4 hours following IRI.

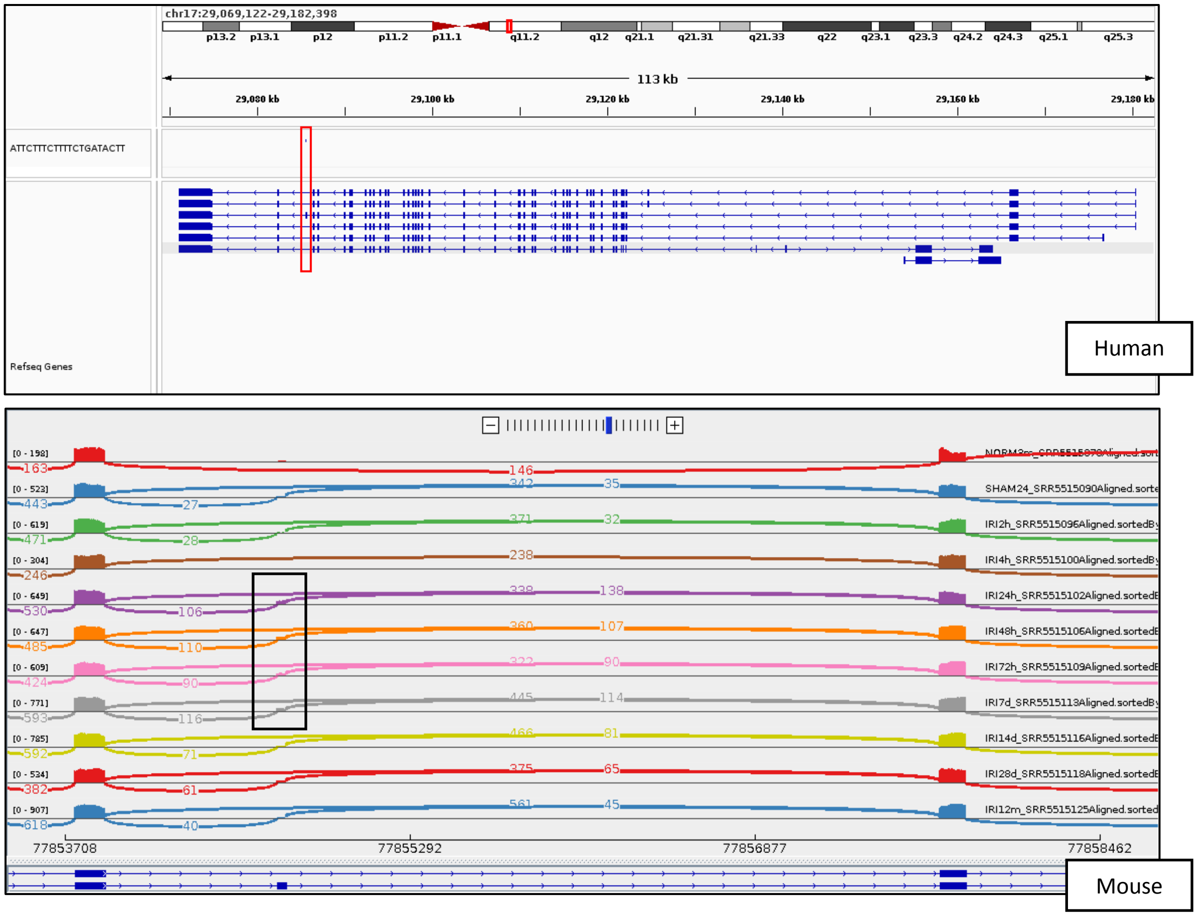


| Time point following IRI | Inclusion left | Inclusion right | inclusion mean | Skipping | Myo18a Exon 48 PSI (approximated) |
|---|---|---|---|---|---|
| 2h | 28 | 32 | 30 | 371 | 0.074812968 |
| 4h | 1 | 1 | 1 | 238 | 0.0041841 |
| 24h | 106 | 138 | 122 | 338 | 0.265217391 |
| 48h | 110 | 107 | 108.5 | 360 | 0.231590181 |
| 72h | 90 | 90 | 90 | 322 | 0.218446602 |
| 7d | 116 | 114 | 115 | 445 | 0.205357143 |
| 14d | 71 | 81 | 76 | 466 | 0.140221402 |
| 28d | 61 | 65 | 63 | 375 | 0.143835616 |
| 12m | 40 | 45 | 42.5 | 561 | 0.070422535 |

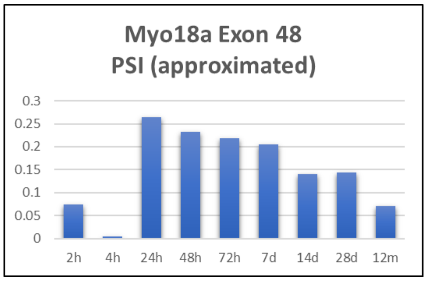


**Figure S9: Exon 48 of the gene Myo18a is somewhat over-expressed from 24 hours to 7 days following IRI.**

Apart from the alternatively spliced exon shown in Figure 2C, we observed that another exon of Myo18a, exon no. 48, is somewhat elevated following IRI. The inclusion of this exon in humans was previously found to be controlled by NEK2 (Figure S11), a non-typical RBP that is also over-expressed following IRI. In humans, NEK2 also controls exon 30 of the gene SPAG9, whose mouse homolog is also over-expressed following IRI in the dataset that we studied (Figure S10). Both exon 48 of MYO18A and exon 30 of SPAG9 were previously found to be associated with EMT in breast cancer (Naro et al. 2021). Top: shown is the siRNA sequence from Naro et al. (Naro et al. 2021) aligned to the alternatively spliced exon 48 in the human MYO18A gene. Bottom: A sashimi plot of selected representative samples (from exon 47 to 49) of the mouse Myo18a showing elevated inclusion levels of exon 48 from 24 hours to 7 days after IRI. Approximated percentage inclusion levels (PSI) were calculated according to junction counts only.

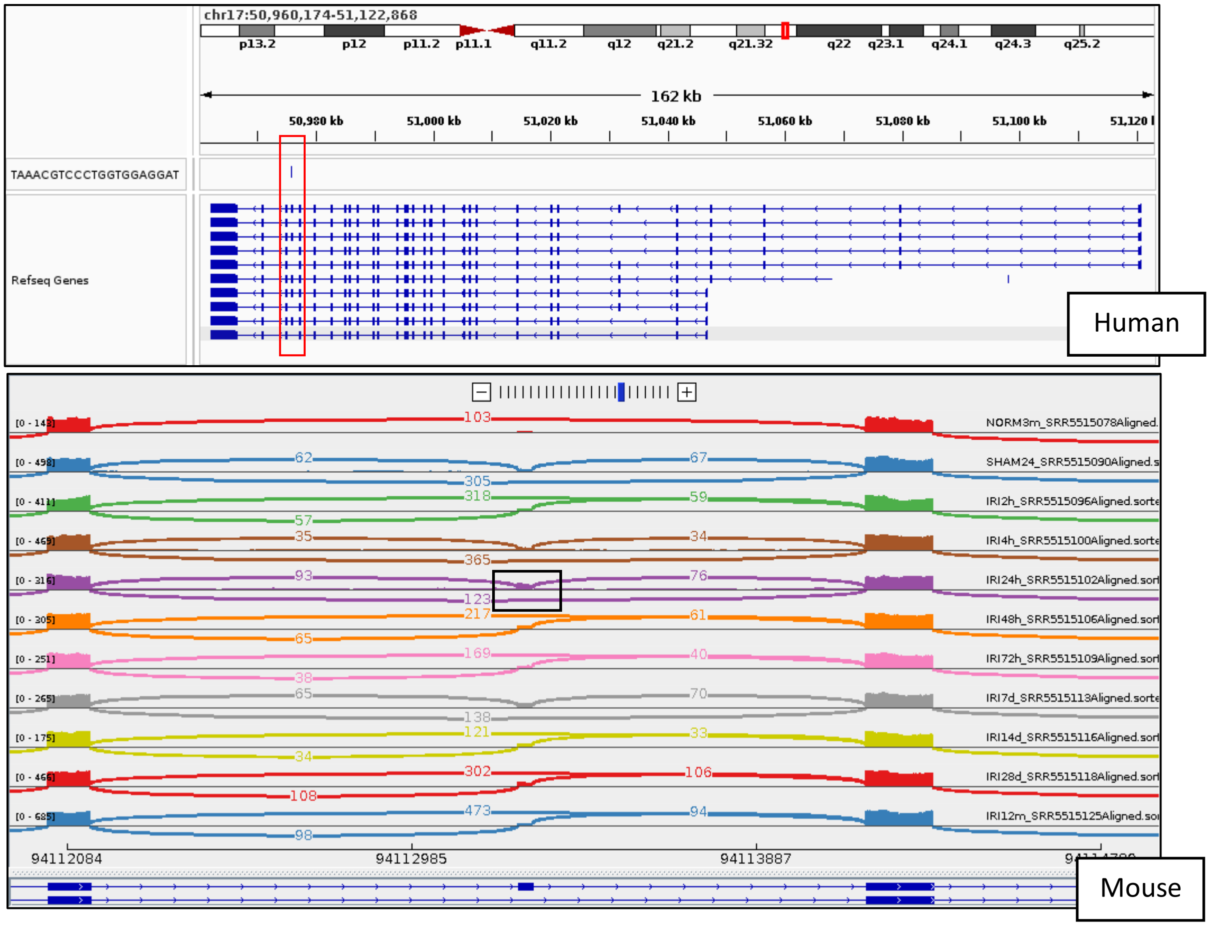


| Time point following IRI | Inclusion left | Inclusion right | inclusion mean | Skipping | Spag9 Exon 30 PSI (approximated) |
|---|---|---|---|---|---|
| 2h | 57 | 59 | 58 | 318 | 0.154255319 |
| 4h | 35 | 34 | 34.5 | 365 | 0.086357947 |
| 24h | 93 | 76 | 84.5 | 123 | 0.407228916 |
| 48h | 65 | 61 | 63 | 217 | 0.225 |
| 72h | 38 | 40 | 39 | 169 | 0.1875 |
| 7d | 65 | 70 | 67.5 | 138 | 0.328467153 |
| 14d | 34 | 33 | 33.5 | 121 | 0.216828479 |
| 28d | 108 | 106 | 107 | 302 | 0.261613692 |
| 12m | 98 | 94 | 96 | 473 | 0.168717047 |

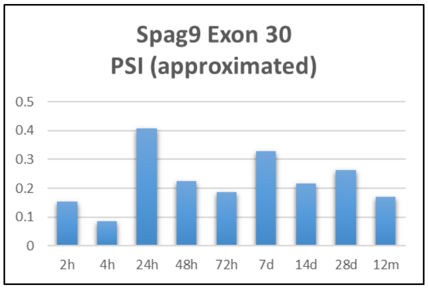


**Figure S10: Exon 30 of the gene Spag9 is somewhat over-expressed from 24 hours following IRI.**

Top: Identification of the alternatively spliced exon. Shown is an siRNA sequence from Naro et al. (Naro et al. 2021) aligned to exon 30 in the human SPAG9 gene. Bottom: A sashimi plot of selected representative samples (from exon 29 to 31) of the mouse Spag9 showing elevated inclusion levels of exon 30 at 24 hours after IRI. Approximated percentage inclusion levels (PSI) were calculated according to junction counts only.

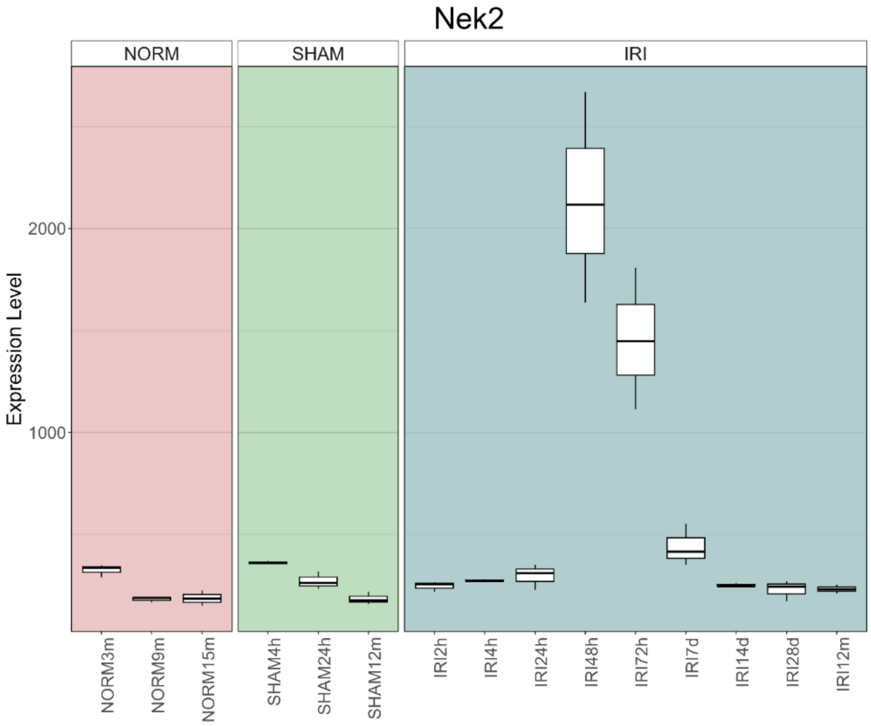


**Figure S11: The RNA binding protein Nek2 is over-expressed from 48 hours to approximately 7 days following IRI.**

Shown is a gene expression boxplot for the gene Nek2, an RNA splicing regulator that is known to promote inclusion of exon 48 of MYO18A and exon 30 of SPAG9 (Naro et al. 2021). Note though that the expression levels of Nek2 rise after the inclusion of the exons that it is known to promote.

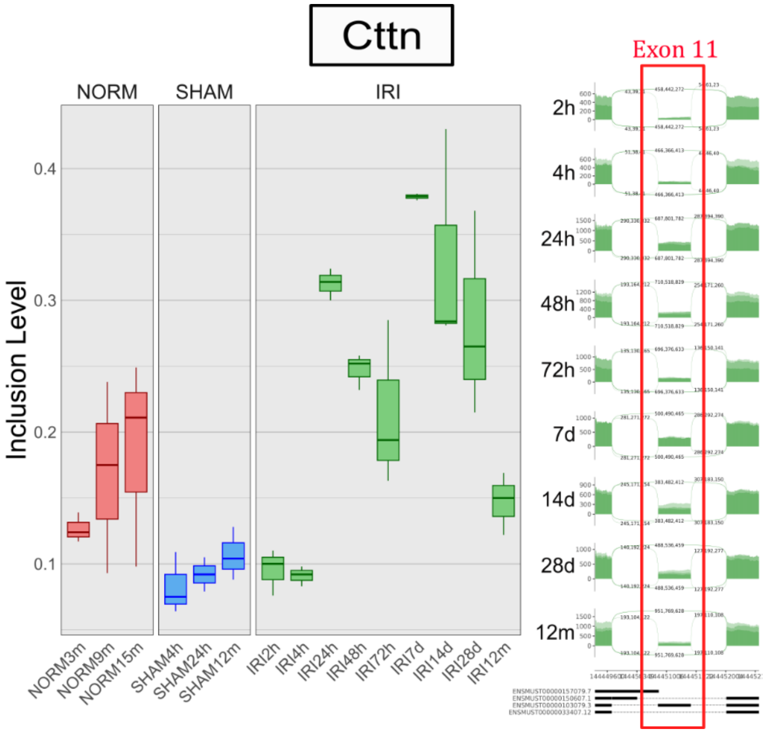


**Figure S12: Exon 11 of the gene Cttn undergoes alternative splicing and is elevated from 24 hours to 28 days following IRI.**

A boxplot of inclusion levels and sashimi plot showing upregulation of exon 11 from 24 hours to 28 days after IRI. Each time-point consists of three replicate samples. Note that the sashimi plot was drawn only for the samples following IRI and not for the normal and sham surgery controls.

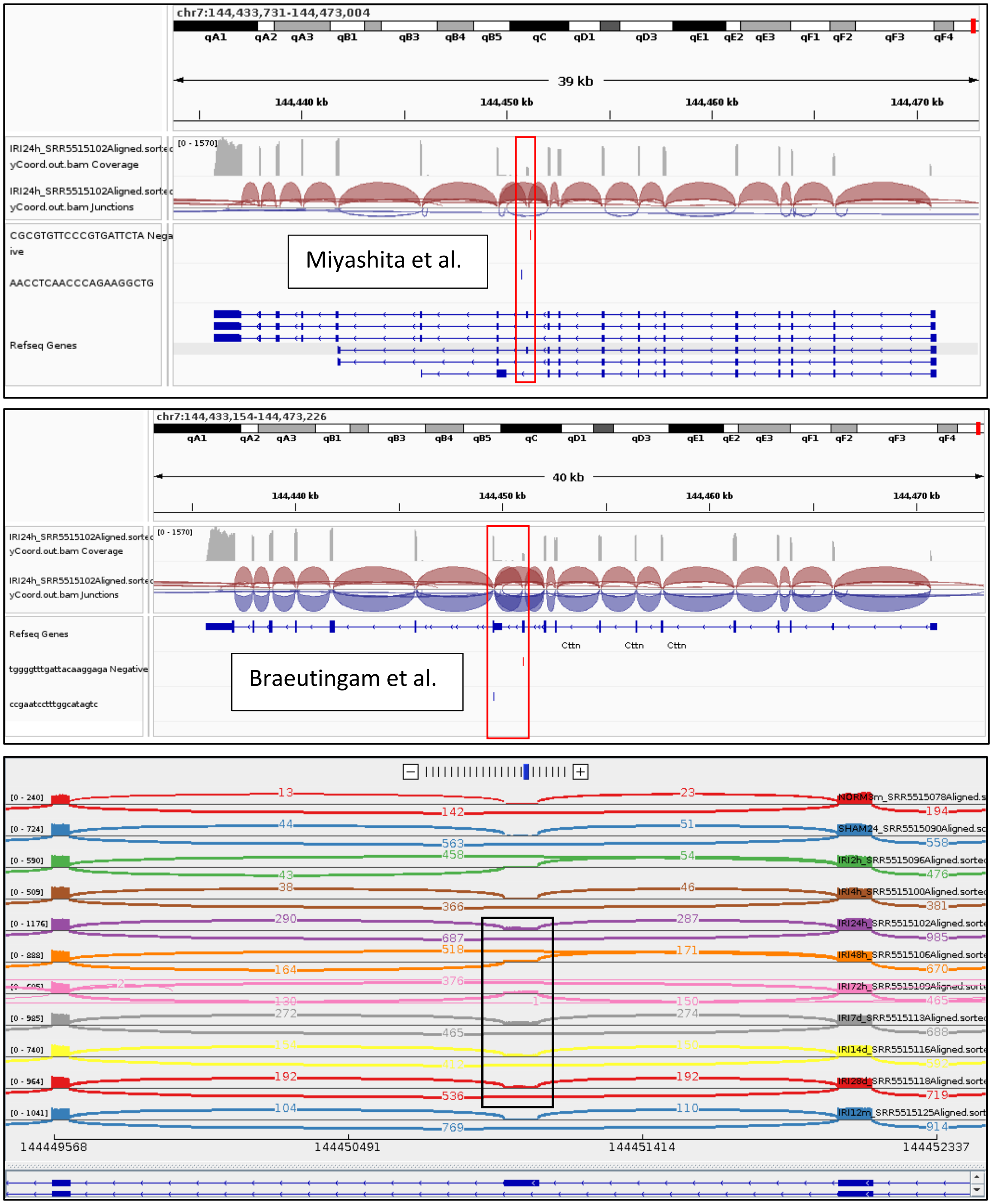


**Figure S13: Exon 11 of the gene Cttn is elevated from 24 hours to 28 days following IRI.**

Top: Identification of this alternatively spliced exon. Shown are PCR primer sequences for exon 11 in the mouse Cttn gene as identified in Miyashita et al. (Miyashita et al. 2021) Middle: Shown are PCR primer sequences for exons 11 and 12 in the mouse Cttn gene as identified in Braeutingam et al. (Braeutigam et al. 2014). Bottom: A sashimi plot of selected representative samples showing elevated inclusion levels of exon 11 from 24 hours to 28 days following IRI.

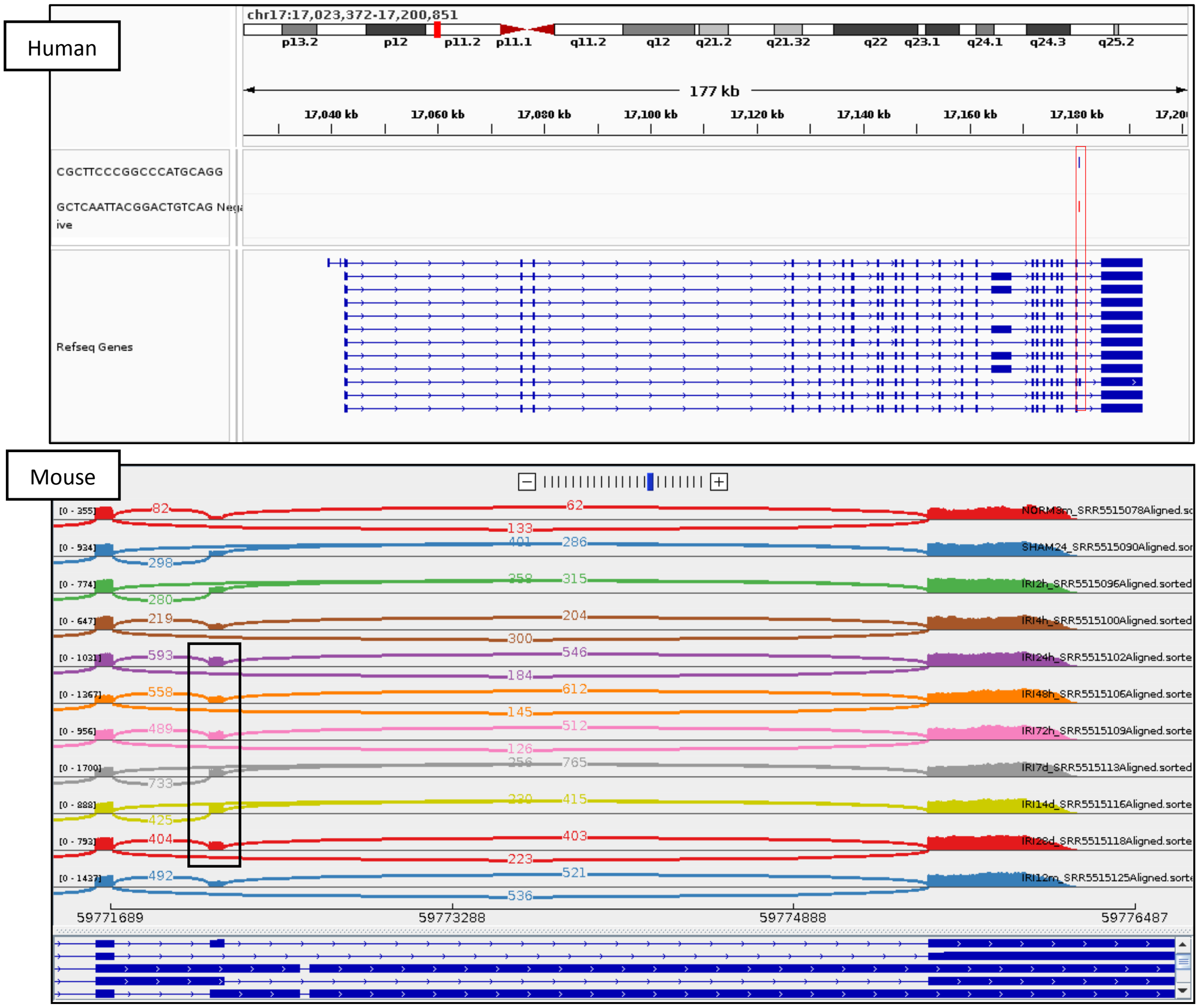


**Figure S14: Exon 23 of the gene Mprip is over-expressed from 24 hours to 28 days following IRI.**

Top: Identification of the alternatively spliced exon. Shown are PCR primer sequences aligned to exon 23 in the human MPRIP gene as identified in Jbara et al. (Jbara et al. 2023). Bottom: A sashimi plot of representative samples of the mouse Mprip gene showing elevated inclusion levels of exon 23 from 24 hours to 28 days after IRI.

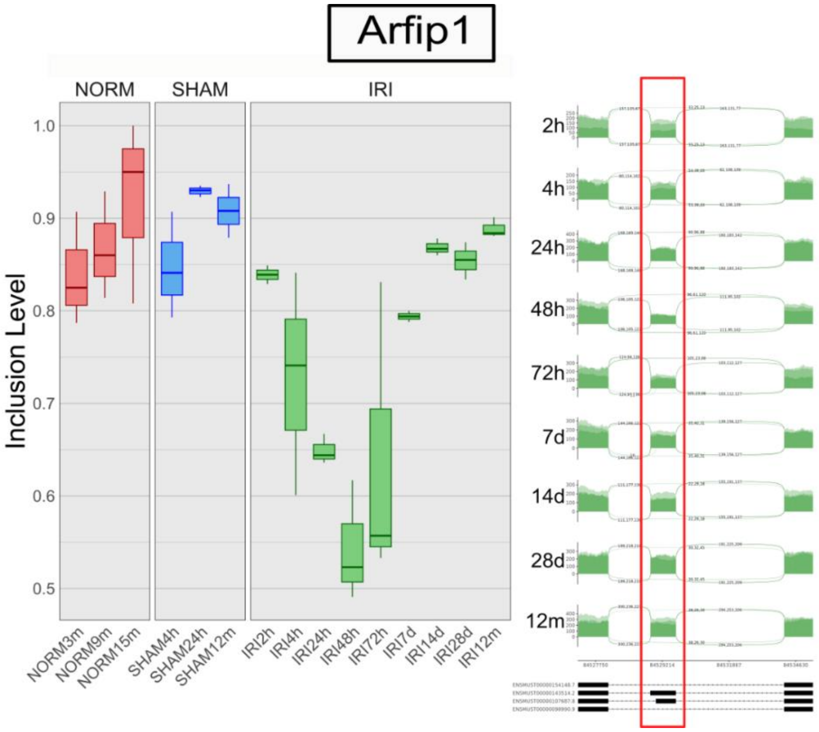


**Figure S15: Arfip1 undergoes alternative splicing following IRI.**

A boxplot of inclusion levels and a sashimi plot showing downregulation of a cassette exon from 4 hours to 7 days after IRI. Each time-point consists of three replicate samples. Note that the sashimi plot was drawn only for the samples following IRI and not for the normal and sham surgery controls.

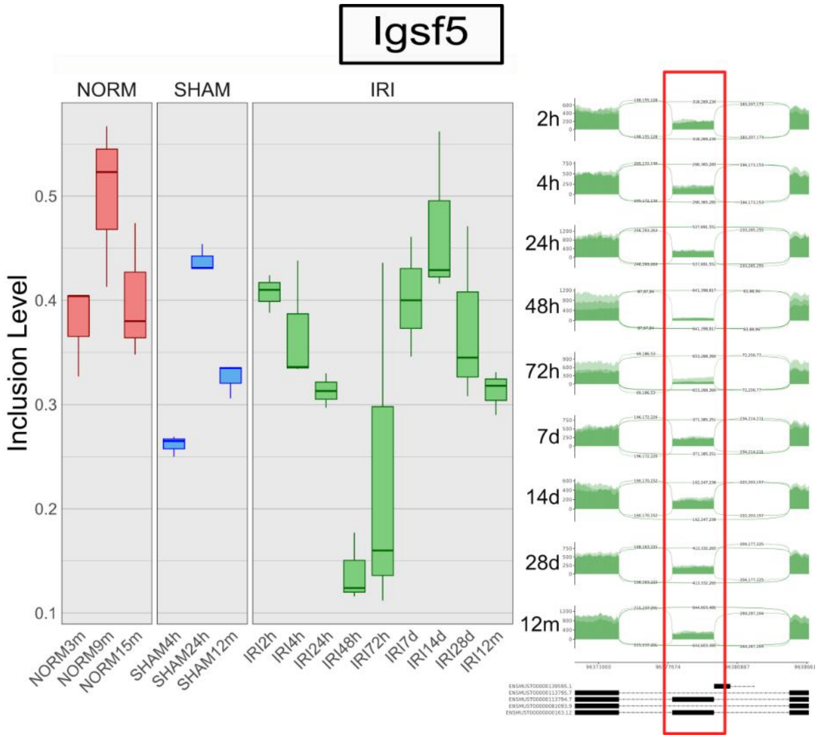


**Figure S16: Igsf5 undergoes alternative splicing following IRI.**

A boxplot of inclusion levels and a sashimi plot showing downregulation of a cassette exon from 48 to 72 hours following IRI. Each time-point consists of three replicate samples. Note that the sashimi plot was drawn only for the samples following IRI and not for the normal and sham surgery controls.

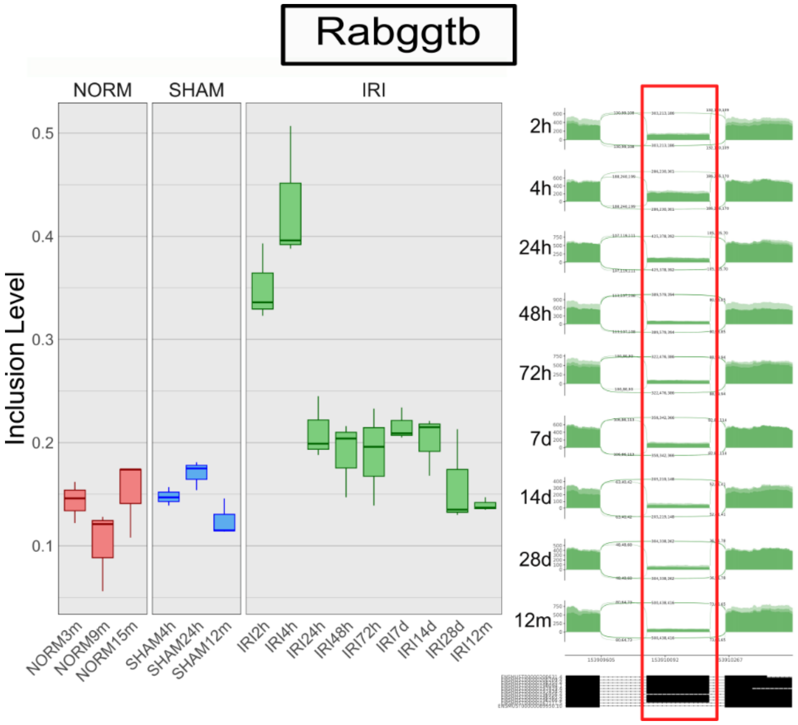


**Figure S17: Rabggtb undergoes alternative splicing following IRI.**

A boxplot of inclusion levels and a sashimi plot showing upregulation of a cassette exon from 2 to 4 hours following IRI. Each time-point consists of three replicate samples. Note that the sashimi plot was drawn only for the samples following IRI and not for the normal and sham surgery controls.

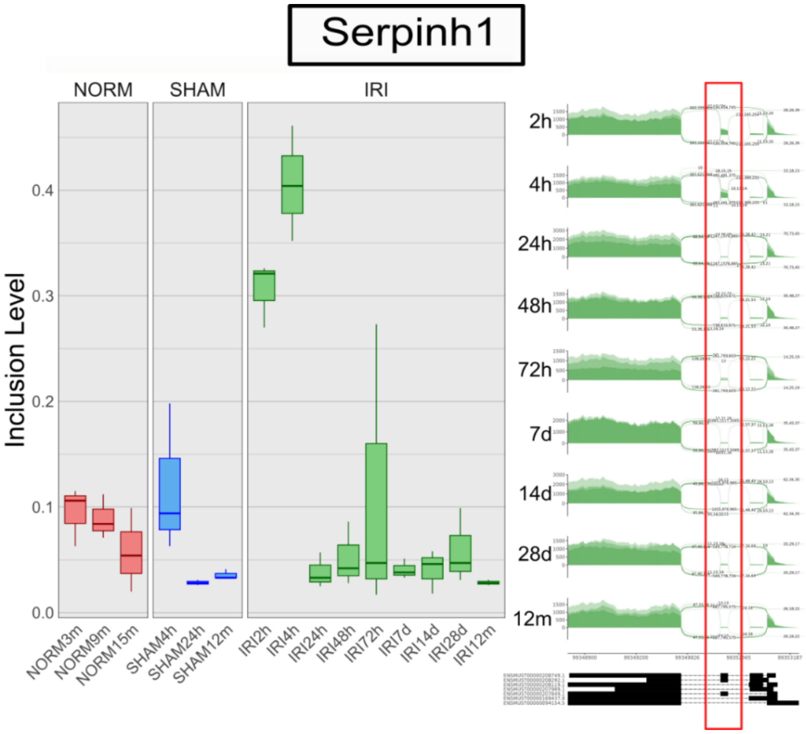


**Figure S18: Serpinh1 undergoes alternative splicing following IRI.**

A boxplot of inclusion levels and a sashimi plot showing upregulation of a cassette exon from 2 to 4 hours following IRI. Each time-point consists of three replicate samples. Note that the sashimi plot was drawn only for the samples following IRI and not for the normal and sham surgery controls.

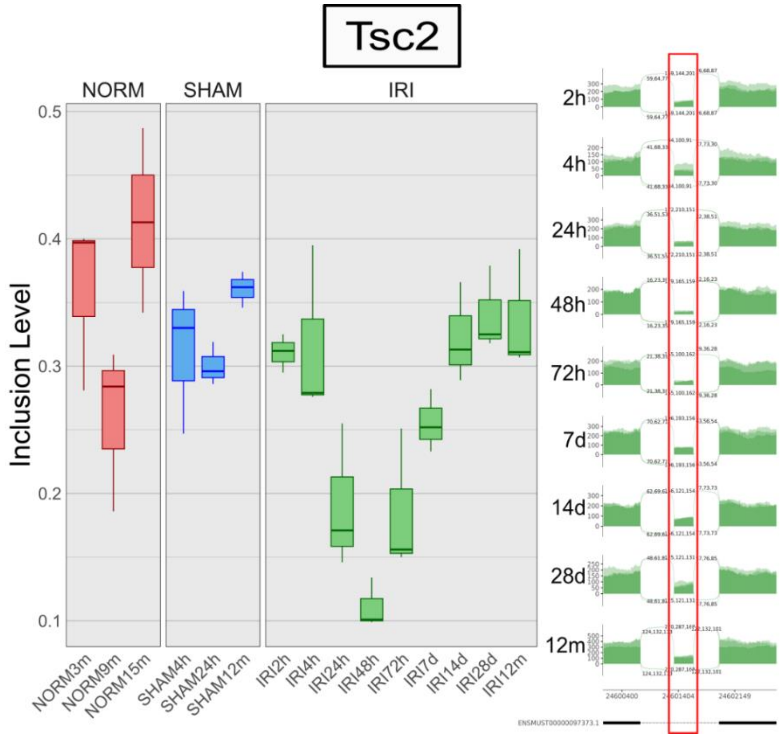


**Figure S19: Tsc2 undergoes alternative splicing following IRI.**

A boxplot of inclusion levels and a sashimi plot showing downregulation of a cassette exon from 24 hours to 7 days following IRI. Each time-point consists of three replicate samples. Note that the sashimi plot was drawn only for the samples following IRI and not for the normal and sham surgery controls.

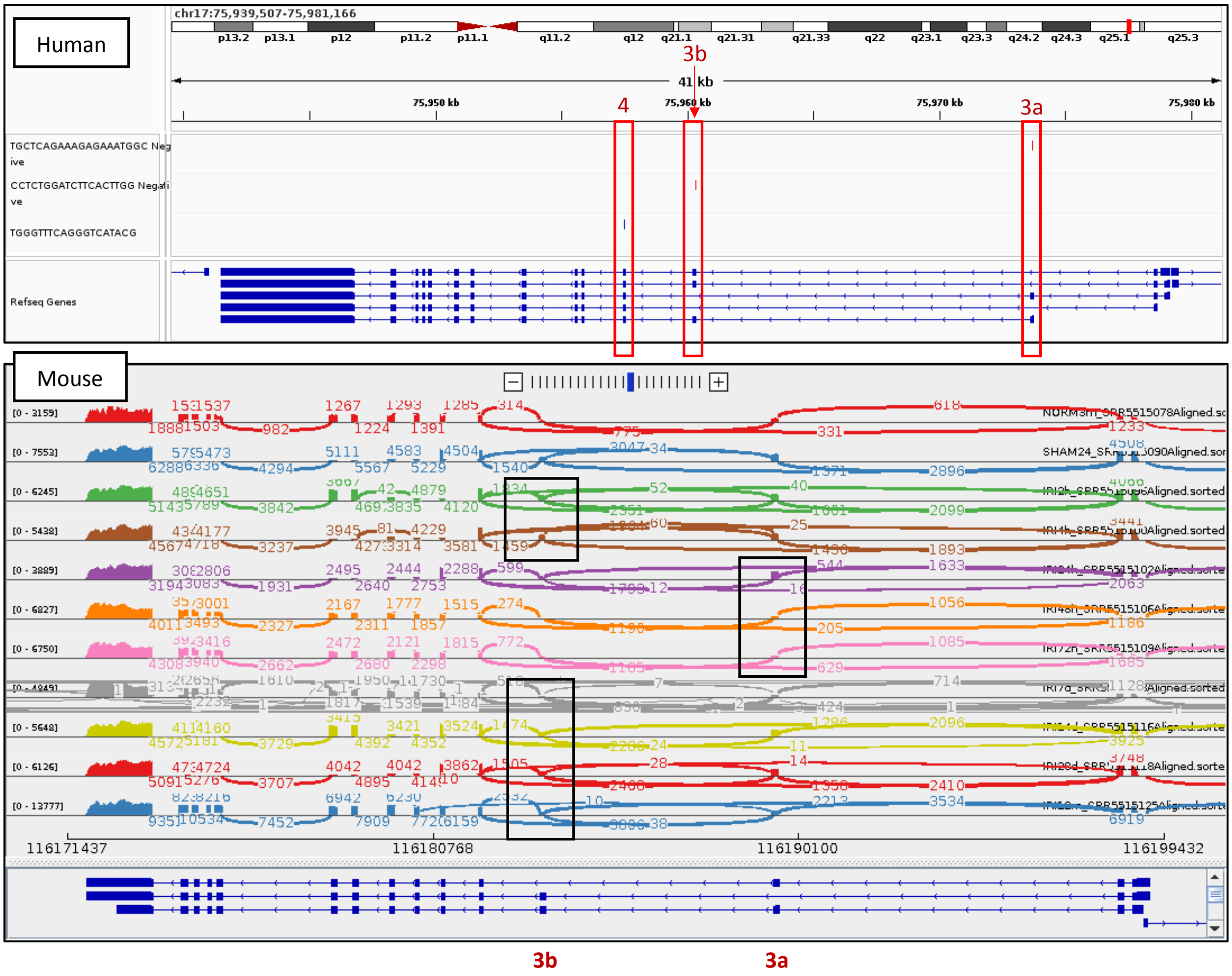


**Figure S20: Exon 3a of the gene Acox1 is over-expressed relative to exon 3b from 24 to 72 hours following IRI.**

Top: Identification of the alternatively spliced exons 3a and 3b. Shown are PCR primer sequences for exons 3a, 3b, and 4 in the human ACOX1 gene as identified in Oaxaca-Castillo et al. (Oaxaca-Castillo et al. 2007). Bottom: A sashimi plot of selected representative samples of the mouse Acox1 gene showing elevated inclusion levels of exon 3a with respect to exon 3b from 24 to 72 hours after IRI.

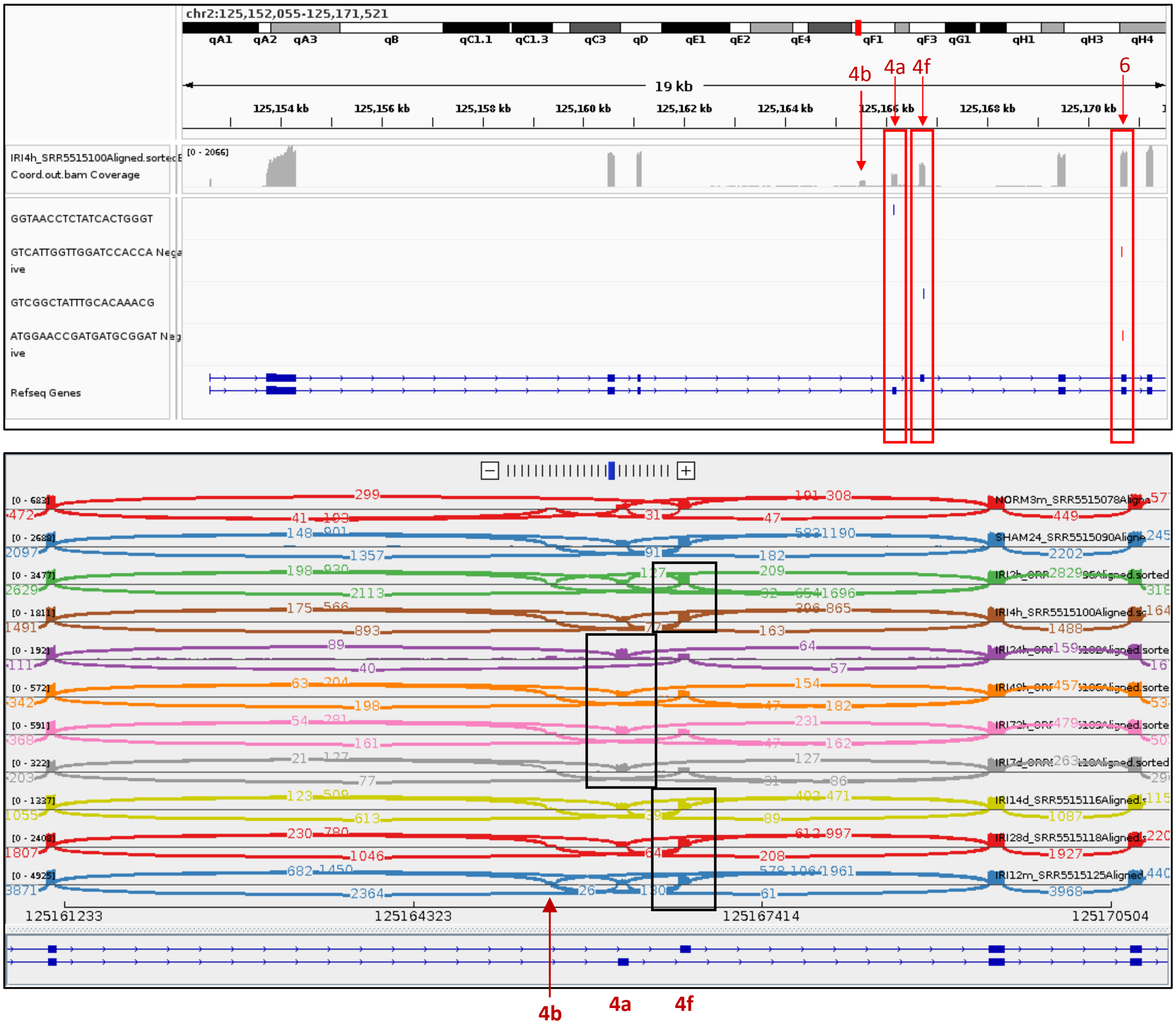


**Figure S21: Exon 4a of the gene Slc12a1 is over-expressed relative to exon 4f from 24 hours to 7 days following IRI.**

Top: Identification of the alternatively spliced exons 4a and 4f. Shown are PCR primer sequences for exons 4a, 4f, and 6 in the mouse Slc12a1 gene as identified in Hao et al. (Hao, Bellner, and Ferreri 2013). Bottom: A sashimi plot of selected representative samples of the mouse Slc12a1 gene showing elevated inclusion levels of exon 4a with respect to exon 4f from 24 hours to 7 days following IRI.

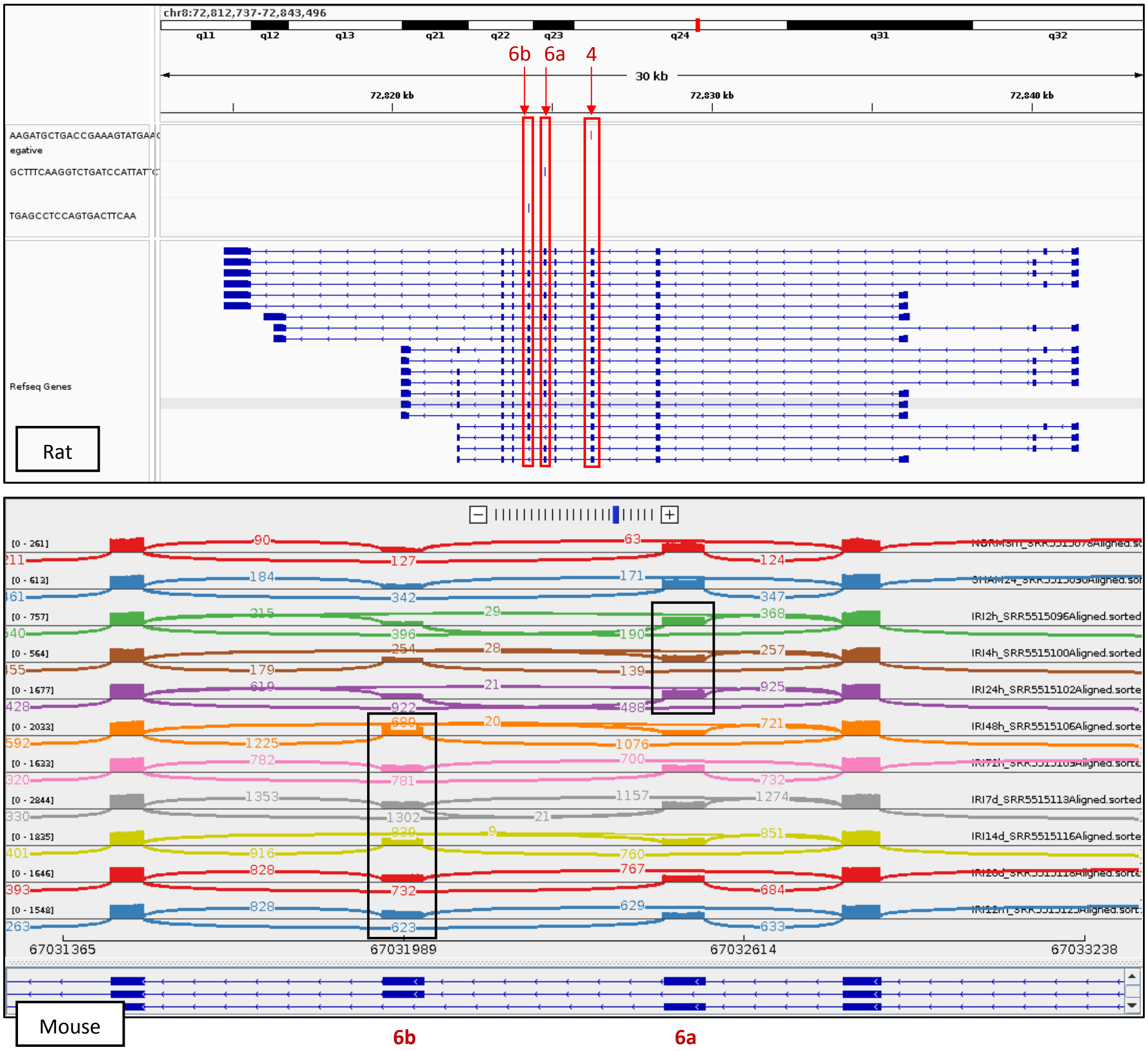


**Figure S22: Exon 6b of the gene Tpm1 is over-expressed relative to exon 6a from 48 hours to 12 months following IRI.**

Top: Identification of the alternatively spliced exons 6a and 6b. Shown are PCR primer sequences for exons 4, 6a, and 6b in the rat Tpm1 gene as identified in Cao et al. (Cao, Routh, and Kuyumcu-Martinez 2021). Bottom: A sashimi plot of selected representative samples of the mouse Tpm1 gene showing elevated inclusion levels of exon 6b with respect to exon 6a from 48 hours to 12 months after IRI.

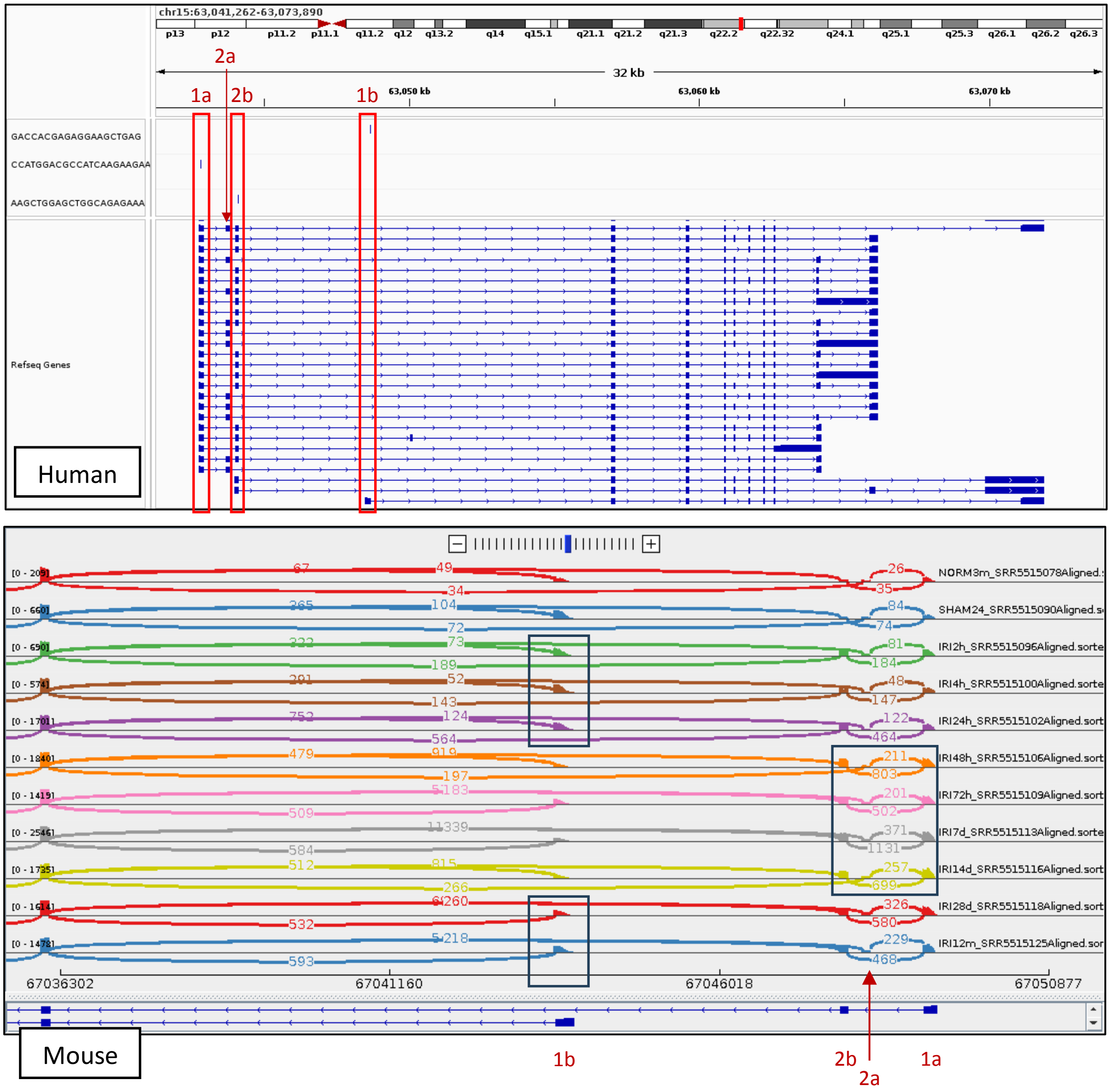


**Figure S23: Exons 1a and 2b of the gene Tpm1 (transcripts "Tpm1.6/7") are over-expressed relative to exon 1b (transcripts "Tpm1.8/9") from 48 hours to 14 days following IRI.**

Top: Identification of the alternatively spliced exons 1a, 2b, and 1b. Shown is an siRNA target sequence for exon 2b and PCR primer sequences for exons 1a and 1b in the human TPM1 gene as identified in Xu et al. (Xu et al. 2024). Bottom: A sashimi plot of selected representative samples of the mouse Tpm1 gene showing elevated inclusion levels of exons 1a and 2b (associated with transcripts "Tpm1.6/7" in Xu et al. (Xu et al. 2024)) with respect to exon 1b (associated with transcripts "Tpm1.8/9") from 48 hours to 14 days after IRI.

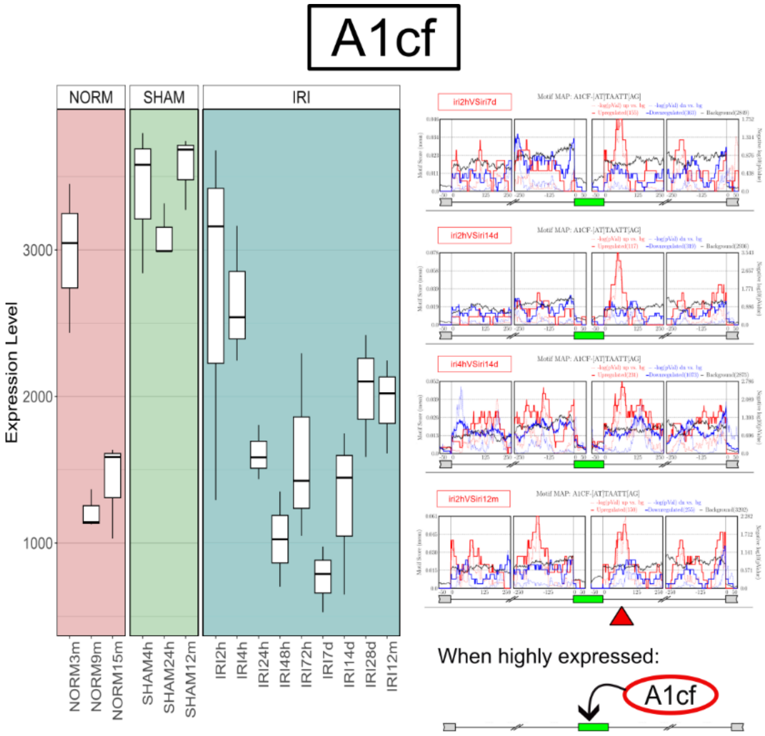


**Figure S24: Motif enrichment analysis indicates that the RNA binding protein A1cf regulates alternative splicing in the kidney following IRI.**

Shown are a gene expression boxplot (left) and motif enrichment diagrams (right) for A1cf. These indicate that A1cf promotes exon inclusion when over-expressed by binding downstream of target exons. The repression of A1cf following IRI is consistent with a previous study by Huang et al. who suggested that knockdown of A1cf in rat normal kidney tubular epithelial cells enhanced EMT by attenuating expression of epithelial markers and inducing mesenchymal ones (Huang et al. 2016).

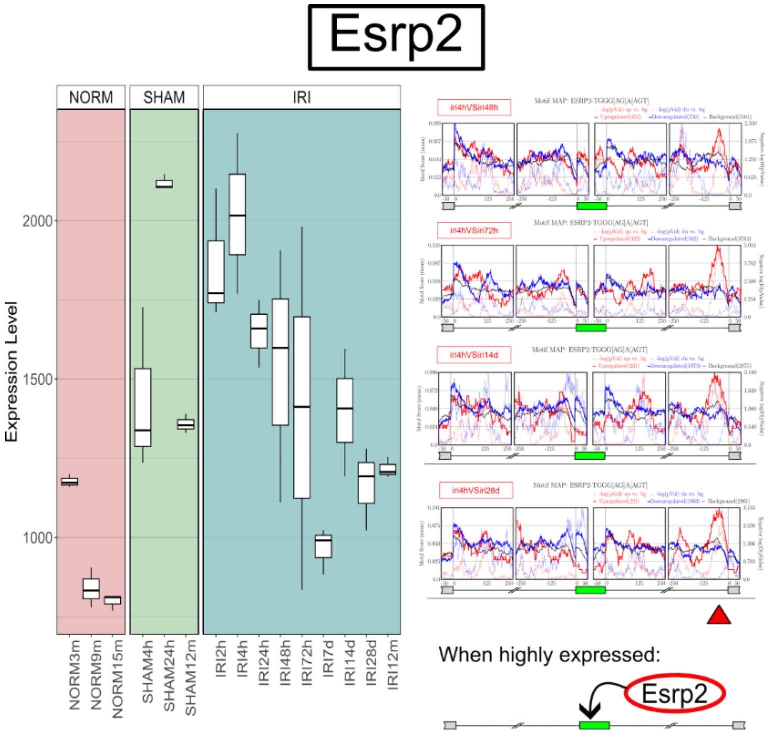


**Figure S25: Motif enrichment analysis indicates that the RNA binding protein Esrp2 regulates alternative splicing in the kidney following IRI.**

Shown are a gene expression boxplot (left) and motif enrichment diagrams (right) for Esrp2. These indicate that Esrp2, which is known to promote the inclusion of epithelial related cassette exons, promotes exon inclusion when over-expressed by binding downstream of target exons.

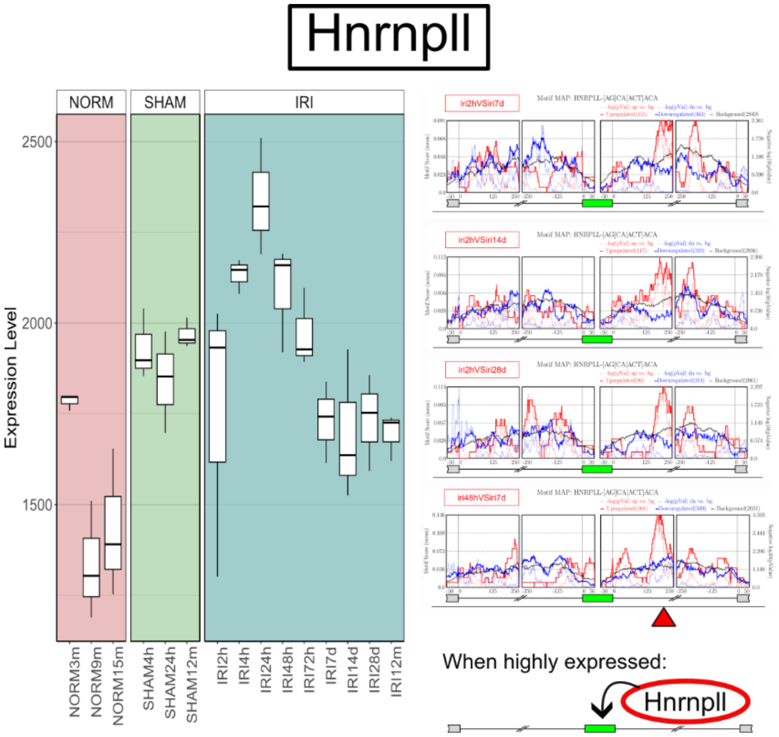


**Figure S26: Motif enrichment analysis indicates that the RNA binding protein Hnrnpll regulates alternative splicing in the kidney following IRI.**

Shown are a gene expression boxplot (left) and motif enrichment diagrams (right) for Hnrnpll. These indicate that Hnrnpll promotes exon inclusion when over-expressed by binding downstream of target exons.

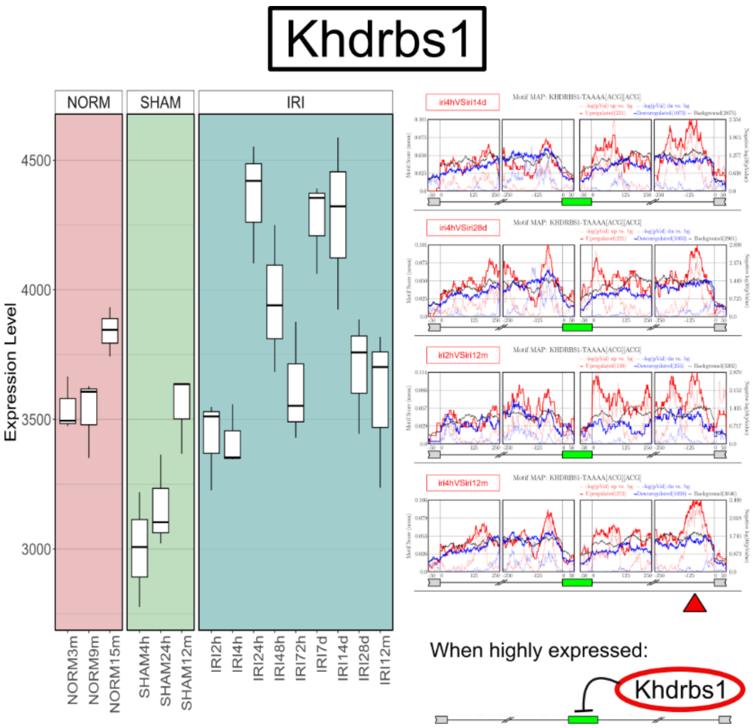


**Figure S27: Motif enrichment analysis indicates that the RNA binding protein Khdrbs1 regulates alternative splicing in the kidney following IRI.**

Shown are a gene expression boxplot (left) and motif enrichment diagrams (right) for Khdrbs1. These indicate that Khdrbs1 represses exon inclusion when over-expressed by binding downstream of target exons.

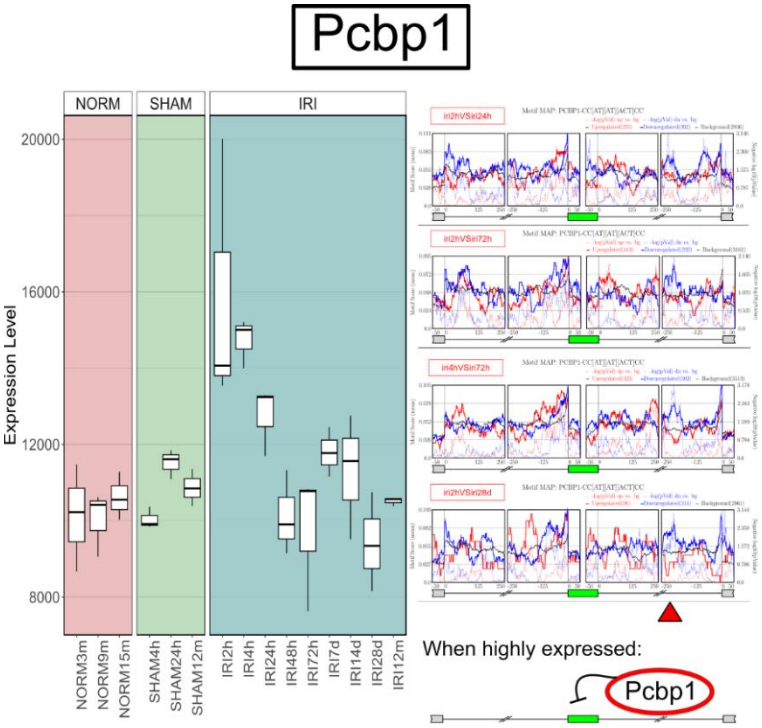


**Figure S28: Motif enrichment analysis indicates that the RNA binding protein Pcbp1 regulates alternative splicing in the kidney following IRI.**

Shown are a gene expression boxplot (left) and motif enrichment diagrams (right) for Pcbp1. These indicate that Pcbp1 represses exon inclusion when over-expressed by binding downstream of target exons. The over-expression of Pcbp1 immediately following IRI is consistent with a previous study where Pcbp1 was found to bind to heavily oxidized RNA to induce apoptosis-related reactions (Ishii et al. 2018).

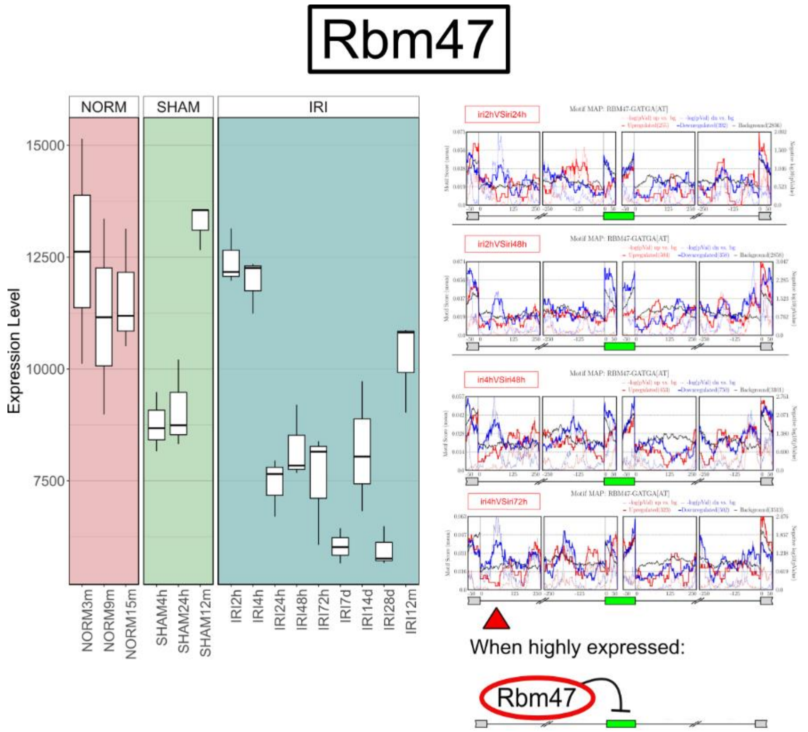


**Figure S29: Motif enrichment analysis indicates that the RNA binding protein Rbm47 regulates alternative splicing in the kidney following IRI.**

Shown are a gene expression boxplot (left) and motif enrichment diagrams (right) for Rbm47. These indicate that Rbm47 represses exon inclusion when over-expressed by binding upstream of target exons.

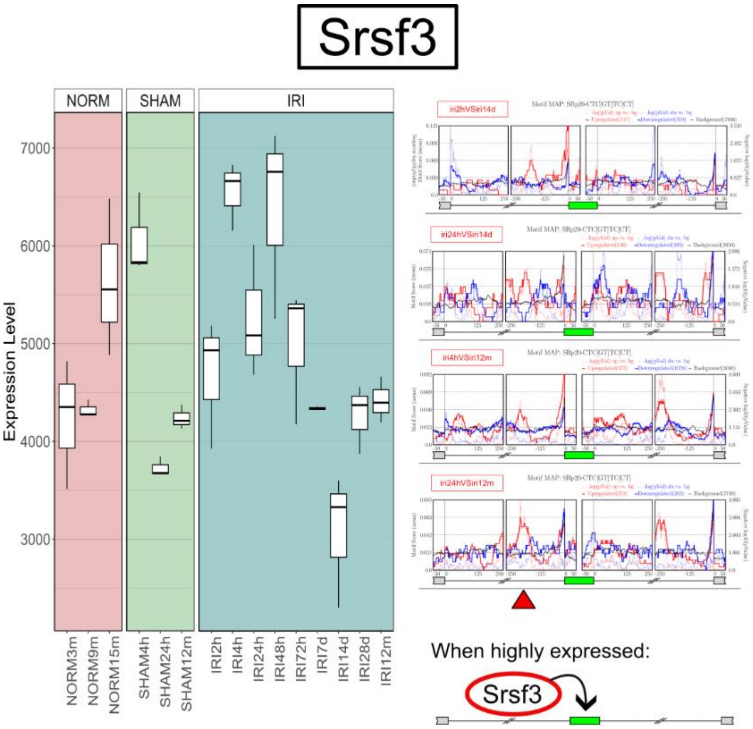


**Figure S30: Motif enrichment analysis indicates that the RNA binding protein Srsf3 regulates alternative splicing in the kidney following IRI.**

Shown are a gene expression boxplot (left) and motif enrichment diagrams (right) for Srsf3. These indicate that Srsf3 promotes exon inclusion when over-expressed by binding upstream of target exons. Note that we found that this gene also undergoes alternative splicing following kidney injury, whereby exon 4 is elevated from 2 to 4 hours following IRI (Figure 2B).

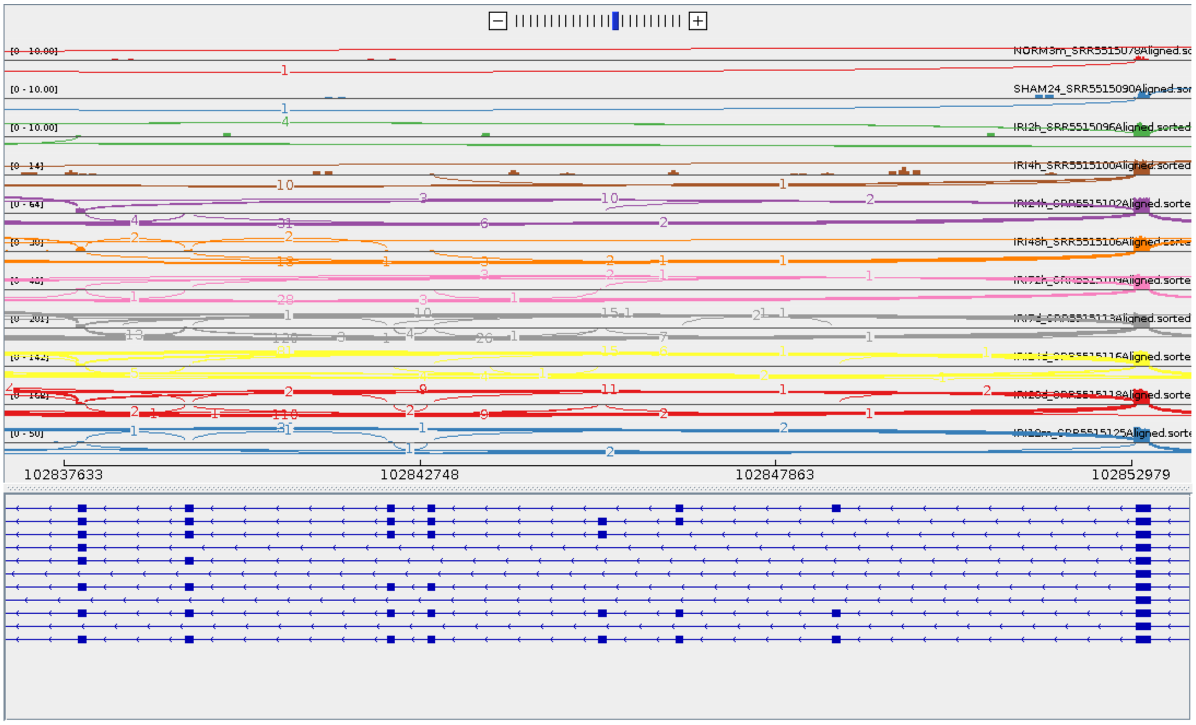


**Figure S31: We did not observe an EMT-related splicing pattern in the gene Cd44.**

Shown is a shashimi plot of the variable exons in Cd44.

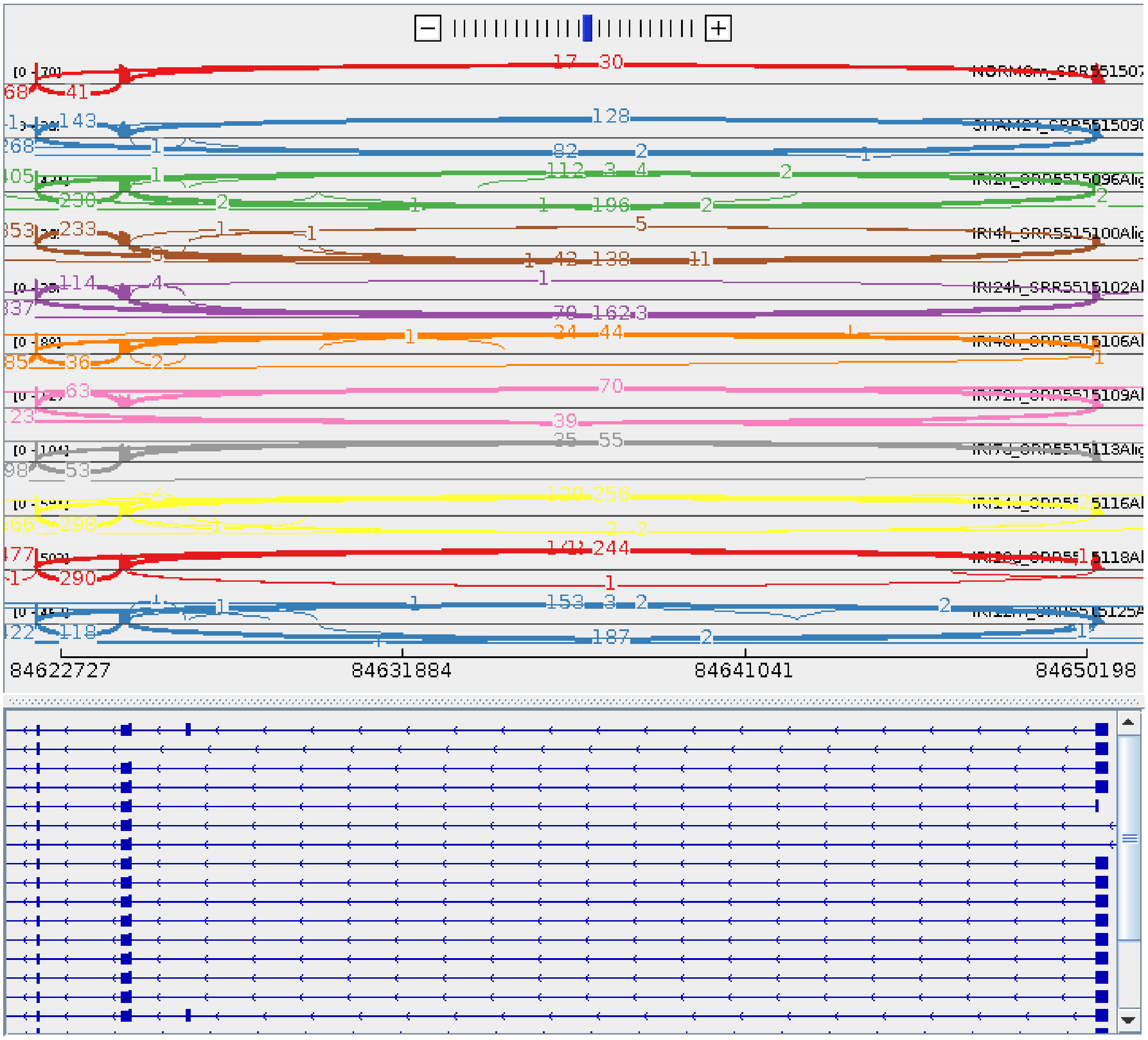

**Figure S32: We did not observe an EMT-related splicing pattern in the gene Ctnnd1.**

Shown is a shashimi plot of the variable exons in Ctnnd1.

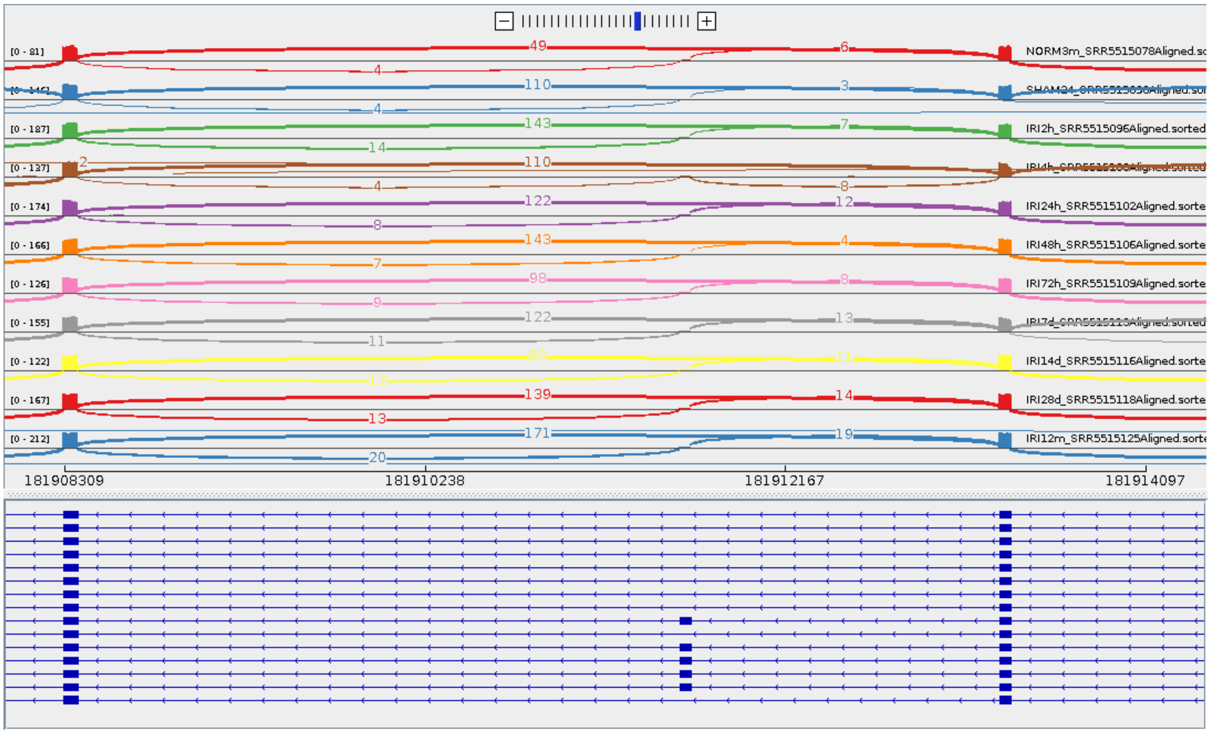


**Figure S33: We did not observe an EMT-related splicing pattern in the gene Enah.**

Shown is a shashimi plot of the variable exon 11a in Enah.

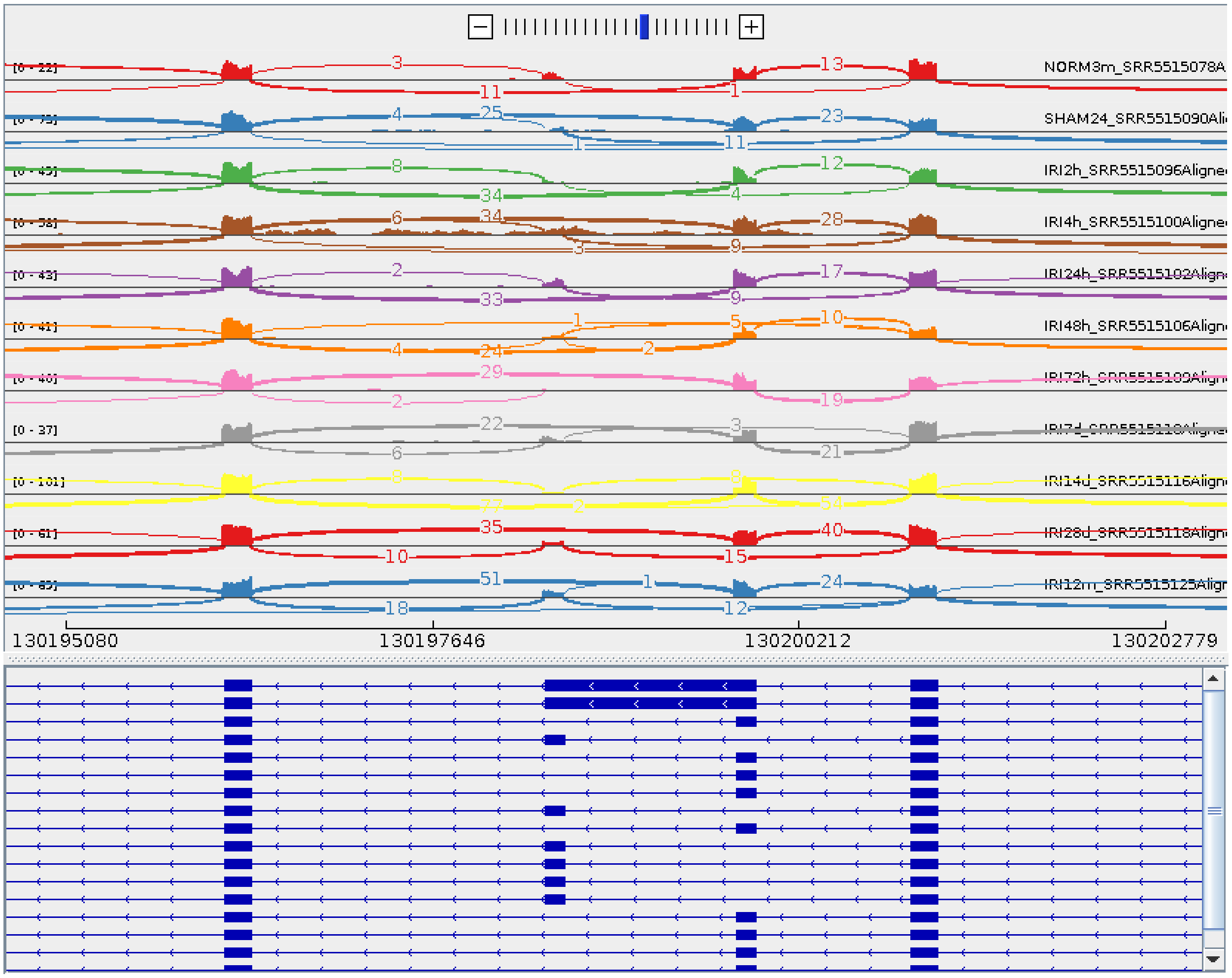


**Figure S34: We did not observe an EMT-related splicing pattern in the gene Fgfr2.**

Shown is a shashimi plot of the mutually exclusive exons in Fgfr2.
